# Dynamic interaction of BcsD subunit in type I bacterial cellulose synthase

**DOI:** 10.1101/2022.03.27.485962

**Authors:** Tatsuya Kondo, Yui Nakamura, Shingo Nojima, Min Yao, Tomoya Imai

**Affiliations:** Research Institute for Sustainable Humanosphere (RISH), Kyoto University, Uji, Kyoto 611-0011, Japan.; Faculty of Advanced Life Science, Hokkaido University, N10W Kita-Ku, Sapporo 060-0810, Japan.

**Keywords:** cellulose, cellulose synthase, membrane enzyme, protein complex, protein-protein interaction, polysaccharide synthesis

## Abstract

Cellulose is a promising biological material for supporting sustainable human life. This natural polymer is synthesized by cellulose synthase, a protein complex in the cell membrane. Cellulose synthase in bacteria is a hetero-subunit complex, and its subunit organization varies widely depending on the species. In the type I bacterial cellulose synthase complex, the BcsD (bacterial cellulose synthase D) protein is believed to play an important role in producing cellulose with long slender fiber morphology and high crystallinity, given the phenotype of the *bcsD*-deficient mutant and the specific existence of the type I operon in bacterial species synthesizing crystalline cellulose microfibrils such as Acetobacter. In this study, we successfully established a heterogeneously expressed Bcs protein in Escherichia coli as an experimental model and conducted biochemical studies for the BcsD protein and the other three major subunits of bacterial cellulose synthase, BcsA, BcsB, and BcsC. It has been shown that the BcsD protein interacts with the functionally required minimal subunits of the BcsAB complex, as well as the BcsC protein. Furthermore, it was shown that BcsD interacts with the BcsAB complex in two modes: direct protein-protein interactions and indirect interactions through the product cellulose. The former and latter modes represent the basal and active states of the type I bacterial cellulose synthase, respectively. This dynamic behavior of the BcsD protein is important for the type I bacterial cellulose synthase complex to regulate the crystallization process of cellulose.

## Introduction

Cellulose is a natural polymer, whose molecular structure is composed of a homopolymer of glucose with a β1→4 linkage, which is very simple. However, cellulose molecules are insoluble in water, and the apparent elementary structure of cellulose is an aggregation of many cellulose molecules in long crystalline fibers. This structural feature of cellulose underlies its structure-sustaining property, which many cellulose-producing organisms rely on.

This polymer assembly is produced by the enzyme “cellulose synthase.” In many known cases, *in vivo* cellulose synthase is a protein complex with several polypeptide subunits. The most important subunit in this complex is glycosyltransferase, which is called CesA. The first structural biological study of the CesA protein was reported in 2013 for cellulose synthase from the bacterium *Rhodobacter sphaeroides* (1, 2), which has stimulated many biochemical studies on cellulose synthase.

Given its simplicity, bacterial cellulose synthase (Bcs) has been a model in many studies on cellulose synthase. Particularly, Bcs from bacteria belonging to the genus *Acetobacter*, some of which are currently named *Gluconacetobacter* or *Komagataeibacter* (the genus name “*Acetobacter*” is used for these bacteria in this study), have been used extensively in studies. Although it is now known that many other bacteria produce cellulose, the function and structure of cellulose appears to vary with species (3). For example, phosphoethanolamine cellulose has been reported in *Escherichia coli*, and this natural cellulose derivative is important for interacting with the host organism (4).

Among cellulose-synthesizing bacteria, *Acetobacter*, which has a gene cluster or operon categorized as type I (3), is characteristic of synthesizing highly crystalline cellulose microfibrils (5) as well as plant cellulose, a major natural source of cellulose. Previous studies have reported that the Bcs-complex in *Acetobacter* is composed of at least four subunits (6, 7) – BcsA (bacterial CesA, catalytic subunit), BcsB (auxiliary subunit having a carbohydrate-binding domain and flavodoxin domain), BcsC (secretion of a glucan chain through the outer membrane), and BcsD (crystallization subunit). In addition, endoglucanase CMCax (8, 9) (glycoside hydrolase family 8 (GH8)), unknown function protein Ccp (8, 10) (also named as BcsH (3)), and β-glucosidase bglAx (GH3) (11) are considered to be functionally involved in the complex. Given that the cooperation of these subunits produces highly crystalline cellulose microfibrils, the spatial organization of these subunits in the cell is important for understanding the cellulose biosynthesis mechanism as a protein function (12).

In these subunits, the BcsD protein is found specifically in bacterial species producing highly crystalline cellulose microfibrils such as *Acetobacter* and *Asaia* (3,13,14), which possess a type I Bcs-operon (3). The *bcsD*-deficient mutant of *Acetobacter* showed a reduction in cellulose production, as well as a lack of ability to synthesize crystalline cellulose microfibrils (6,7,15). Sunagawa *et al.* also demonstrated that BcsD interacts with Ccp (BcsH) in a pull-down assay (16). Given that the BcsD protein contains glucan chains inside its octamer structure (15), it has been proposed that BcsD is functionally located downstream of the BcsAB complex (the cellulose-polymerizing core of the Bcs-complex (1,6,17,18)) (19, 20). Detailed information on how these functional units work together is key to understanding the mechanism by which cellulose synthase can bundle the synthesized glucan chains for spinning a long slender crystalline fiber at room temperature in an aqueous environment.

In this study, we analyzed protein-protein interactions in the type I Bcs-complex using the recombinant BcsABCD protein with the *E. coli* expression system and clarified the interaction of BcsD with BcsAB complex as well as BcsC protein. One of the advantages of using recombinant proteins is that mutant proteins are easily available (21). Owing to this advantage, this study sheds light on the interaction between BcsAB and BcsD and demonstrates that there are two modes for this interaction: direct protein-protein interactions and indirect interactions through the cellulose molecules. We hypothesized that such bimodal interactions between BcsAB and BcsD proteins could be key to controlling cellulose synthesis dynamically *in vivo*.

## Results

### Production of endogenous bcsA gene-deficient E. coli

The expression of the *Acetobacter* Bcs complex in *E. coli* cells might lead to contamination by endogenous *E. coli* Bcs (22).We then attempted to delete *bcsA* in the *E. coli* host, which encodes the catalytic subunit of Bcs. The deletion of the *bcsA* gene in *E. coli* BL21, a strain expressing the Bcs recombinant protein in this study, was performed using the Red/ET recombination system (Gene Bridges GmdH, Germany), in which the *bcsA* gene was replaced by the FRT-PGK-gb2-neo-FRT cassette with homology arms of the *bcsA*-flanking region using homologous recombination (Fig. 1A). This genetically modified BL21 (BL21-GM) strain was confirmed by colony direct polymerase chain reaction (PCR) to contain the neomycin-resistance gene cassette (FRT-PGK-gb2-neo-FRT cassette) (Fig. 1B). When amplified with primer #2 (antibiotic gene cassette-specific) and #3 (*E. coli*-specific), a band of approximately 1,500 bp was found in BL21-GM, while no specific product band was found in wild-type BL21, indicating the insertion of the neomycin-resistance gene cassette specifically in BL21-GM. However, when amplified with primers #1 and #3 (either *E. coli*-specific), wild-type BL21 and BL21-GM produced a band corresponding to the *bcsA* gene (2,818 bp) and the inserted gene cassette (1,731 bp), respectively. These results confirmed the successful deletion of the *bcsA* gene of BL21 in “BL21-GM.”

**Figure 1.**
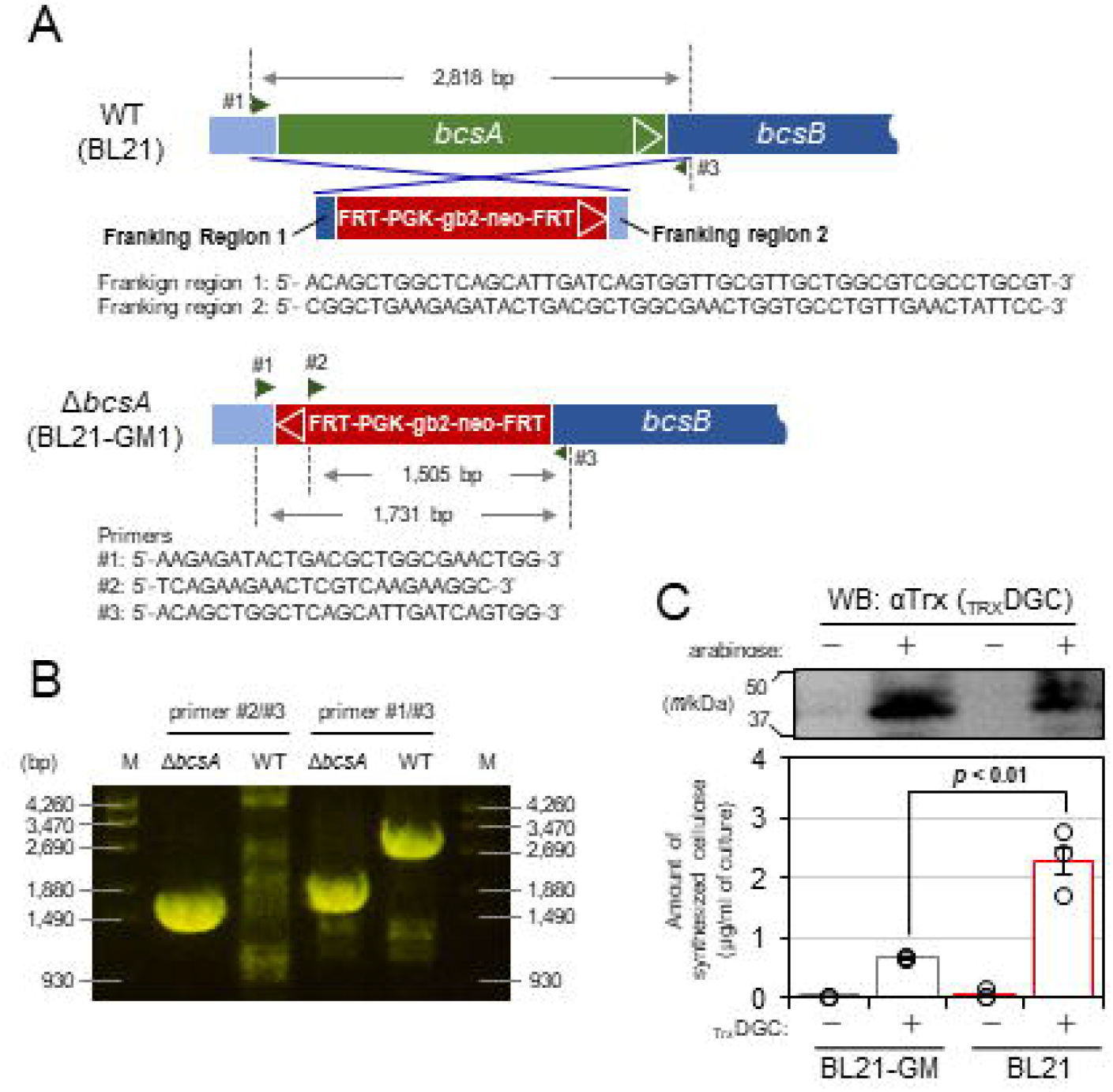
Deletion of the E. coli endogenous bcsA gene. **A** Schematic diagram of the experiment for deleting the endogenous *bcsA* gene using the Red/ET recombination system. The primer sequences used are listed in Table S1. **B** Electrophoresis pattern of colony-direct PCR using two pairs of primers. M: Molecular weight marker (Marker 6 (Nippon Gene Co. Ltd., Tokyo, Japan)); Δ*bcsA*: *bcsA*-deleted BL21; WT: wild-type BL21. **C** Quantification of endogenous cellulose synthesis in BL21-GM and BL21 cells. The CESEC system (18), which uses DGC expression to synthesize c-di-GMP to activate bacterial cellulose synthase, was used to count the amount of cellulose that used DGC expression (bottom). The expression of DGC was confirmed by WB using an anti-Trx antibody (top). The experiments were performed in triplicate (*n* = 3) and are represented as mean ± standard error of the mean (SEM), together with each of the data values. Statistical tests (Student’s t-test) were performed using the built-in function in Microsoft Excel (TTEST).

Next, we measured the *in vivo* cellulose synthesis activity of the prepared BL21-GM (Fig. 1C) by partially using our previous method (18): diguanylate cyclase (DGC) fused to thioredoxin (_Trx_DGC) was expressed in BL21 and BL21-GM to synthesize c-di-GMP to activate endogeneous bacterial cellulose synthase. The amounts of cellulose synthesized were approximately 2.3 and 0.7 μg/mL of culture in BL21 and BL21-GM, respectively. Although weak cellulose-synthesizing activity was observed, we used BL21-GM as a suitable host to express the Bcs subunits with minimal impact on the endogenous Bcs-complex of *E. coli*.

### Characterization of BsCel5A

Cellulase is an important tool used in this study to scrutinize the role of cellulose molecules in the subunit-subunit interaction of the Bcs-complex. In this study, we used a bacterial endo-β-1,4-glucanase from *Bacillus subtilis* (BsCel5A), whose structure has been previously reported (23). The full length (FL) of this cellulase is a two-domain protein, in which the catalytic domain of GH5 is linked to the carbohydrate binding module of family 3 (CBM3). In this study, the catalytic domain alone (CD) was also used to maximize access to the soluble substrate. FL and CD of BsCel5A were expressed as glutathione-*S*-transferase (GST)-fused proteins and purified with glutathione affinity resin, as confirmed by SDS-PAGE (Fig. S1A, left panel); (Fig. S1A, left panel, white arrow). Activity-deficient mutant of E169Q/E257Q in FL (BsCel5A_FLm) and CD (BsCel5A_CDm) was also used for a negative control protein in the cellulase treatment experiment.

The binding activity of the purified BsCel5A to phosphoric acid-swollen cellulose (PASC) was surveyed to confirm correct folding of the expressed proteins. The results showed that all the BsCel5A proteins used in this study retained their binding activity to PASC (Fig. S1A, right panel), indicating that all of them were properly folded despite sequence modification. The enzymatic activity of BsCel5A was surveyed to determine whether this cellulase could function at low temperature and neutral pH, which is required for analyzing the Bcs-complex. Both BsCel5A_FL and BsCel5A_CD hydrolyzed barley β-glucan, carboxyl methyl cellulose (CMC), and PASC at 4℃ and pH 7.5 (Fig. S1B). For Avicel, full-length BsCel5A showed higher activity than the others, stressing the function of CBM to bind solid cellulose.

Substrate specificity of BsCel5A was investigated using cello-oligosaccharides with various degrees of polymerization (DP). BsCel5A hydrolyzed cellotetraose, cellopentaose, and cellohexaose but not cellobiose or cellotriose (Fig. S1C). The final reaction products of BsCel5A for substrates longer than triose were cellobiose and cellotriose, and the smallest substrate was cellotetraose (Fig. S1D).

Summarizing these, both BsCel5A_FL and BsCel5A_CD were able to hydrolyze cellulose of low crystallinity at 4℃ and pH 7.5, and cellulose molecules with DP higher than 3 were the substrates. This information was considered when interpreting the data.

### *In vivo* cellulose synthase activity of recombinant Bcs-complex

This study analyzed heterogeneously expressed Bcs proteins in *E. coli* as an experimental model. We then checked if these heterogeneously expressed proteins were functional by using CESEC (cellulose-synthesizing *E. coli*), a platform for estimating the cellulose-synthesizing activity with the co-expression of diguanylate cyclase (DGC) (18). All the Bcs-complexes expressed in DGC-expressing BL21-GM synthesized a significant amount of cellulose (Fig. 2A), indicating that each of the Bcs-complexes expressed in this study are functional, regardless of whether BcsD is tagged. In addition, neither BcsAB nor the BcsABCD complex with the activity-deficient mutant of the catalytic subunit BcsA (D333N or W373A) showed cellulose-synthesizing activity, consistent with our previous study (21). These results support the idea that the Bcs proteins expressed in this study reflect the state in function and are suitable for an experimental model.

**Figure 2.**
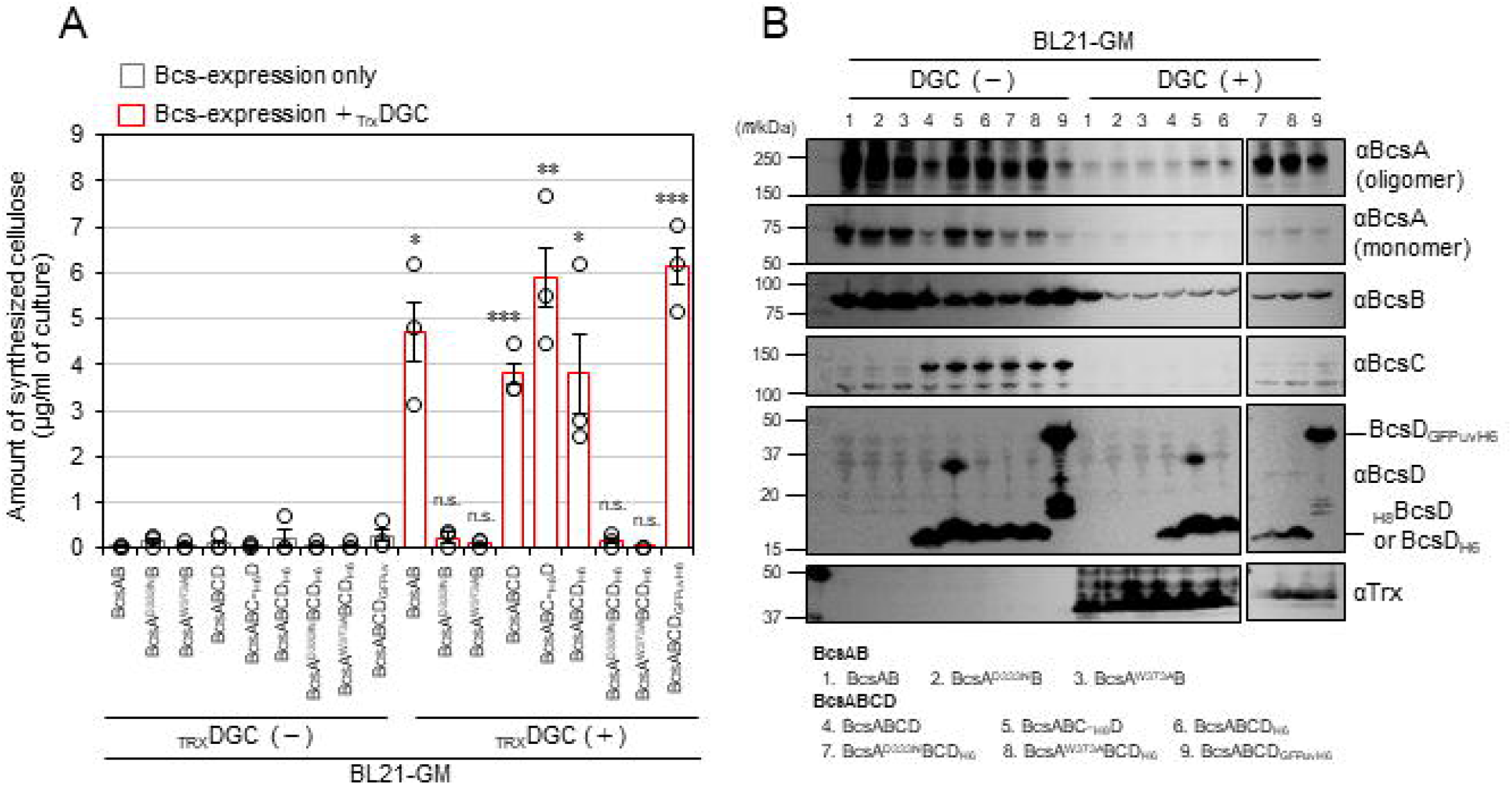
in vivo cellulose-synthesizing activity of recombinant Bcs-complex. **A** Cellulose-synthesizing activity of the recombinant Bcs-complexes tested in this study: the amount of cellulose synthesized by the CESEC system (18) with BL21-GM (Δ*bcsA*) is shown for each recombinant protein. All experiments were performed in triplicate (*n* = 3) and are represented as mean ± SEM together with the significance of difference: * for *p* < 0.05, ** for *p* < 0.01, *** for *p* < 0.001, and *n.s.* for not significant (*p >* 0.1) as a result of Student’s t-test, which was performed using the built-in function of Microsoft Excel (TTEST). **B** Western blotting (WB) was performed for the CESEC experiment to check the expression of BcsA, BcsB, BcsC, BcsD, and _TRX_DGC. Cultures with and without DGC expression were tested. For BcsA, the data are shown separately for the oligomer and monomer bands. Due to the limited number of wells in the SDS-PAGE gel, the gel was divided into two parts for electrophoresis.

Despite the lower cellulose production shown in Fig. 2A, we decided to express Bcs proteins without co-expression of DGC for subsequent purification because higher expression levels of Bcs proteins were detected by western blotting (WB), as shown in Fig. 2B. Given that the cellulose quantification in this study used strong acid treatment with acetic acid and nitric acid, we think that highly crystalline cellulose was majorly considered in the data presented in Fig. 2A. Then it cannot be ruled out that the Bcs-complex expressed in *E. coli* without DGC expression synthesizes cellulose.

### The complex formation of BcsAB, BcsC, and BcsD

Given that BcsD holds the glucan chains in its channel (15), it is functionally located downstream of the BcsAB complex, the core of cellulose molecule synthesis, indicating that BcsD operates in the periplasmic space to assemble cellulose molecules extruded from the BcsAB complex. Furthermore, the BcsC protein has a β-barrel porin-like structure at its C-terminus and is supposed to secrete cellulose through the outer membrane (24). The results from several studies suggest that BcsAB, BscC, and BcsD work together in a coordinated manner to spin bacterial cellulose. We tried to co-express BcsABCD to purify the whole Bcs-complex using a hexa-histidine tag attached to the C-terminus of BcsD (BcsD_H6_), as the tag at the C-terminus does not disturb cellulose-synthesizing activity (16). After the optimization of the conditions (Fig. S2), all four subunits were successfully purified together by immobilized metal affinity chromatography (IMAC) (Fig. 3). This is the first successful purification of the BcsABCD complex ever reported. However, BcsC was found to be mostly insoluble in detergent (*n*-dodecyl-β-D-maltoside, DDM, in this study) and tended to degrade or aggregate during the purification process (Fig. 3A, BcsC antibody strip). We did not decipher BcsC behavior in this Bcs-complex purification experiment.

**Figure 3.**
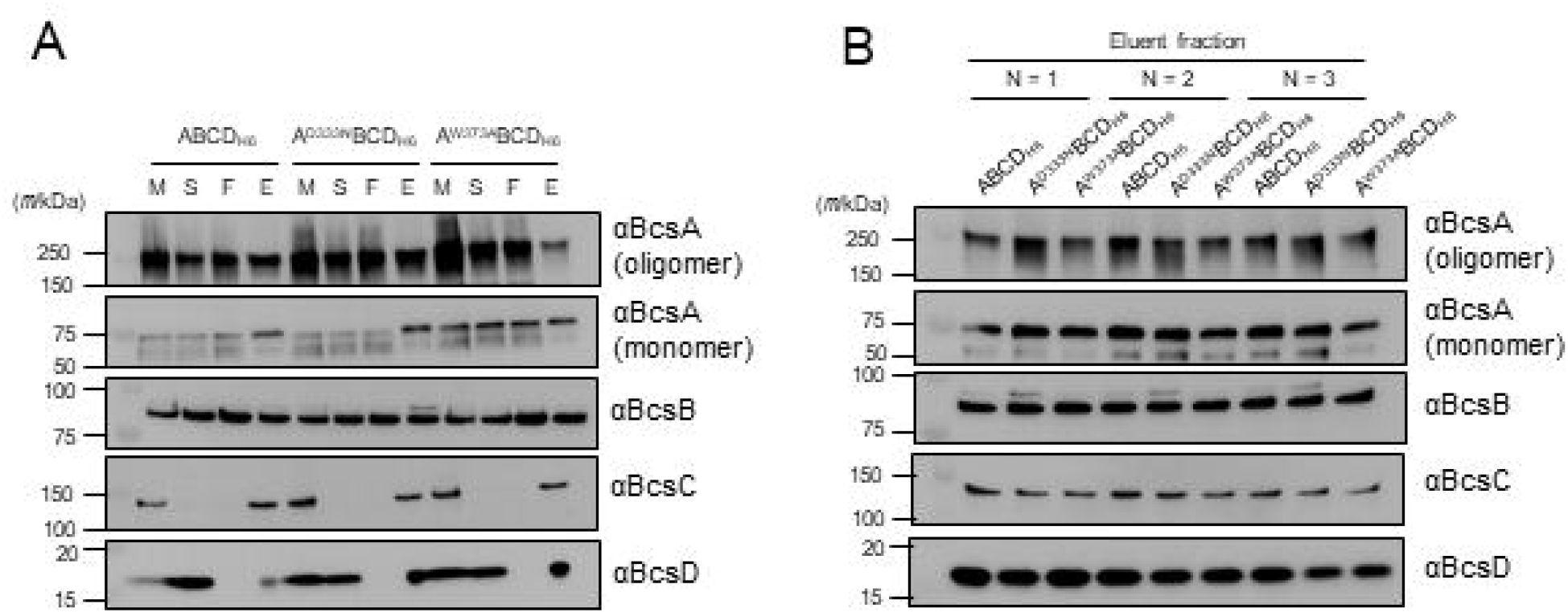
Purification of BcsABCD_H6_. **A** Western blotting (WB) analysis of the purification process: M, membrane fraction; S, solubilized fraction; F, IMAC unbound fraction; E, IMAC-eluted fraction. Three protein samples, wild-type (BcsABCD_H6_), D333N on BcsA (BcsA^D333N^BCD_H6_), and W373A on BcsA (BcsA^W373A^BCD_H6_), were analyzed using the antibodies indicated. **B** WB analyses of the eluate from three independent purification experiments for each of the three protein samples. Approximately 2 µg of the protein was loaded onto the gel for WB with BcsA, BcsB, and BcsC antibodies, and 1 µg of the protein was added to the gel for WB with BcsD antibody. For BcsA, the data are shown separately for the oligomer and monomer bands.

### Analysis of the interaction between BcsAB and BcsD by cellulase treatment

As described above, co-expression of BcsABCD_H6_ in *E. coli* allowed the BcsABD-complex to be purified. Given that BcsABCD_H6_ expressed in *E. coli* is functional as shown in Fig. 2A, we speculated that a glucan chain may link BcsAB and BcsD during this purification. This hypothesis was tested by checking whether the digestion of cellulose resulted in the dissociation of BcsAB from BcsD_H6_ during IMAC purification. BsCel5A cellulase was applied to BcsABCD_H6_ bound to IMAC resin prior to washing and elution. The amounts of BcsA, BcsB, and BcsD proteins in the eluate were measured by semi-quantitative WB (Fig. 4A); the reliability of quantification in our WB was confirmed, as shown in Fig. S3. The signal intensities of BcsA and BcsB were normalized to that of BcsD in the eluate for comparison between samples. Notably, the amounts of BcsA and BcsB bound to BcsD were decreased by BsCel5A treatment (Fig. 4B), indicating that the BcsAB complex was detached from BcsD by cellulose digestion. This trend was clearer when BsCel5A_CD was used as cellulase.

**Figure 4.**
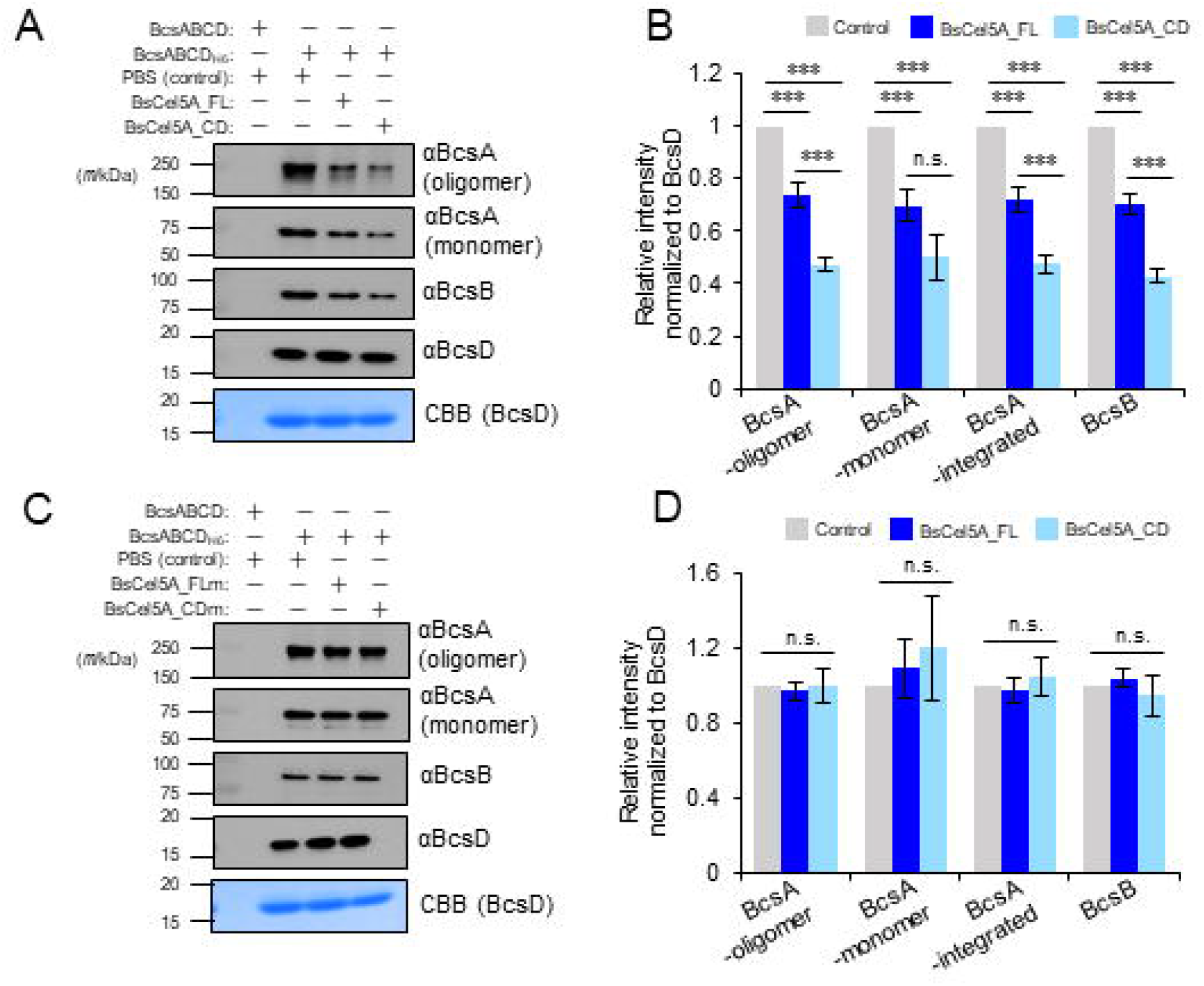
Effect of cellulase treatment to the purification of Bcs-complex. **A**, Western blot images of IMAC-eluted fractions of BcsABCD_H6_ after treatment with wild-type cellulase (BsCel5A_FL and BsCel5A_CD). Coomassie Brilliant Blue (CBB)-stained images were used as loading controls. **B** Band intensity in western blot analysis of *A*. Error bars indicate SEM, and the significance of the difference as a result of Student’s *t*-test, which was performed using the built-in function of Microsoft Excel (TTEST), is indicated by the symbols: ** for *p* < 0.01, *** for *p* < 0.001, and *n.s.* for not significant (*p* > 0.1). **C**, Western blot images of IMAC-eluted fractions of BcsABCD_H6_ after cellulase mutant (BsCel5A_FLm and BsCel5A_CDm) treatment. CBB-stained images were visualized using BcsD as a loading control. **D**, Band intensity in western blot analysis of *C*. Error bars indicate SEM, and Student’s *t*-test, which was performed using the built-in function of Microsoft Excel (TTEST), showed no significant difference (*p >* 0.1; eight independent replicates). For BcsA, the data are shown separately for the oligomer band, monomer band, and their sum. Each elution sample (∼0.2 mg/mL) was added to an SDS-PAGE gel in 10 µL (anti-BcsA and anti-BcsB) and 1 µL (anti-BcsD and CBB stain), respectively. Elution samples for CBB staining were loaded into gel wells 10-fold more than those for WB (BcsD lane).

To rule out the possibility that the observed detachment of BcsAB was due to non-specific interaction of the added BsCel5A protein, the activity-deficient mutant of BsCel5A (E169Q/E257Q) was applied to BcsABCD_H6_ bound to the metal affinity resin (Fig. 4, C and D). As a result of purification, no decrease in the signal intensity of BcsA and BcsB was observed in the eluate, indicating that cellulose hydrolysis actually caused detachment of BcsAB from BcsD during purification. In summary, the BcsAB complex indirectly interacts with BcsD through the glucan chain.

Although cellulase treatment allowed the BcsAB complex to be detached from BcsD during purification, as demonstrated above, a substantial portion of the BcsA and BcsB proteins remained with BcsD in this purification process (Fig. 4B). We supposed that a direct interaction between BcsAB and BcsD is also found in addition to the indirect interaction. To test this hypothesis, we attempted to purify the activity-deficient mutants of BcsABCD_H6_, BcsA^D333N^BCD_H6_, and BcsA^W373A^BCD_H6_, neither of which synthesize cellulose, as hown in Fig. 2A. As a result, these mutants allowed BcsA and BcsB to be purified together with BcsD and the wild type (Fig. 5A and B). Notably, cellulase treatment did not reduce the amount of BcsAB purified together with BcsD for either BcsA^D333N^BCD_H6_ or BcsA^W373A^BCD_H6_ (Fig. 5C, 5D, and 5E). These results support the existence of a direct interaction between BcsAB and BcsD when cellulose is not synthesized. Compiling these cellulase treatment experiments, it was demonstrated that there are two modes of interaction between BcsAB and BcsD: direct and indirect modes through cellulose molecules.

**Figure 5.**
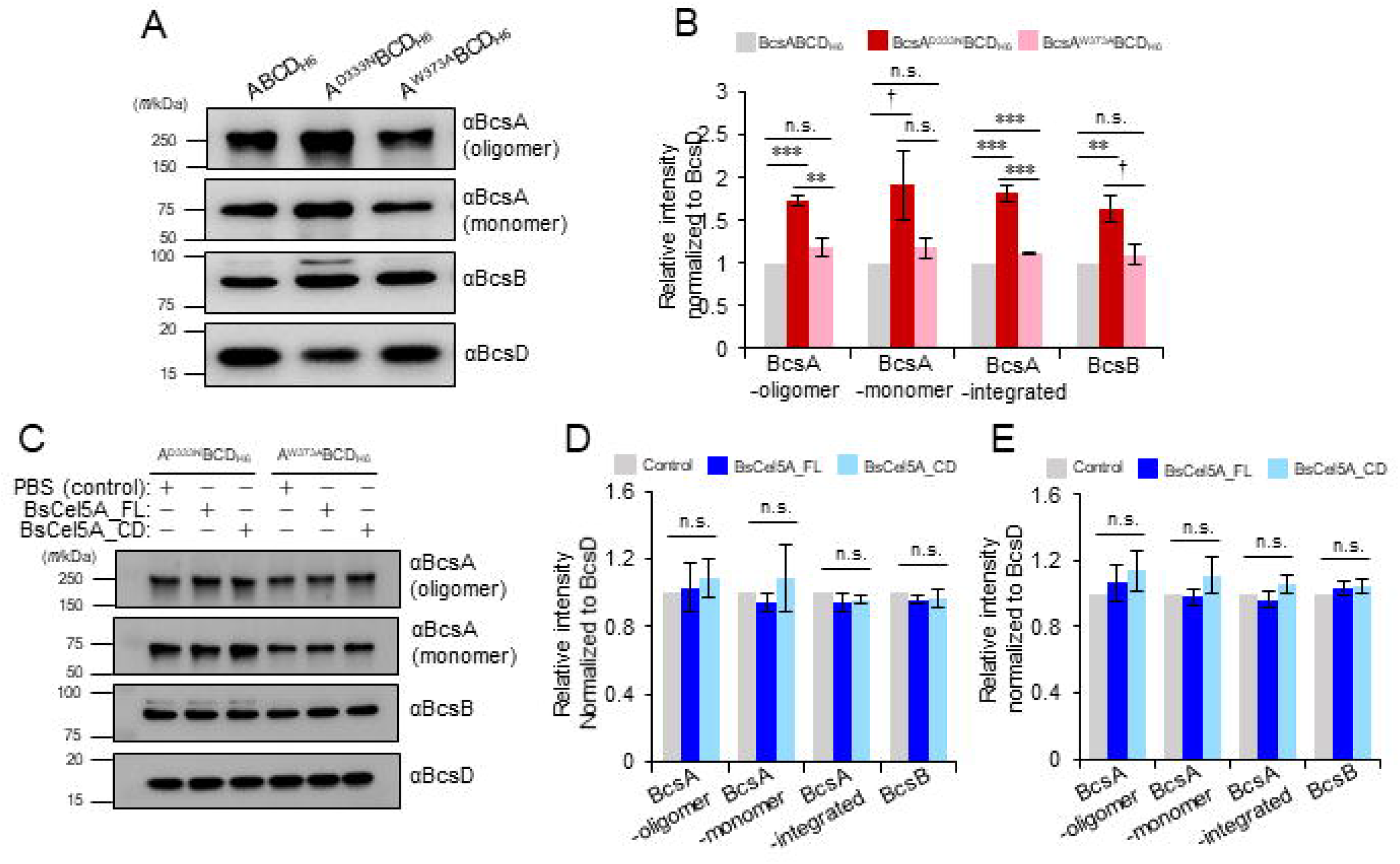
Effect of cellulase treatment on the purification of mutant Bcs-complex. **A** Western blot images of IMAC-eluted fractions of BcsABCD_H6_, BcsA^D333N^BCD_H6_, and BcsA^W373A^BCD_H6_. CBB-stained images were visualized using BcsD as a loading control. **B** Band intensity in western blot analysis of *A*. An error bar indicates SEM, and the significance of the difference as a result of Student’s *t*-test, which was performed using the built-in function of Microsoft Excel (TTEST), is indicated by the symbols † for *p* < 0.1, ** for *p* < 0.01, *** for *p* < 0.001, and *n.s.* for not significant (*p* > 0.1). **C**, Western blot images of IMAC-eluted fractions of BcsA^D333N^BCD_H6_ and BcsA^W373A^BCD_H6_ after treatment with wild-type cellulase (BsCel5A_FL and BsCel5A_CD). **D** and **E** Band intensity on western blot analysis of **C**: D for BcsA^D333N^BCD and E for BcsA^W373A^BCD. An error bar indicates SEM, and the significance of the difference as a result of Student’s *t*-test, which was performed using the built-in function of Microsoft Excel (TTEST), showed no significant difference between any dataset. All experiments were performed with four independent replicates (*n* = 4). For BcsA, the data are shown separately for the oligomer band, monomer band, and their sum. Each elution sample (∼0.2 mg/mL) was added to an SDS-PAGE gel in 10 µL (anti-BcsA and anti-BcsB) and 1 µL (anti-BcsD and CBB stain), respectively. Elution samples for CBB staining were loaded into gel wells 10-fold more than those for WB (BcsD lane).

### Quaternary structure of the Bcs-complex

Despite the demonstration of two modes for the interaction between BcsAB and BcsD, the quaternary structure of the entire Bcs-complex remains to be elucidated. We then performed pull-down assays to further explore the subunit-subunit interactions in the BcsABCD complex by fully utilizing recombinant Bcs proteins. The assay was designed based on the results from previous studies (Fig. 6A): (i) BcsAB forms a complex in the inner membrane (1), (ii) BcsC is an outer membrane protein (24, 25), (iii) BcsD protein is located in the periplasmic space (15, 20), and (iv) BcsD interacts with the BcsAB complex (this study).

**Figure 6.**
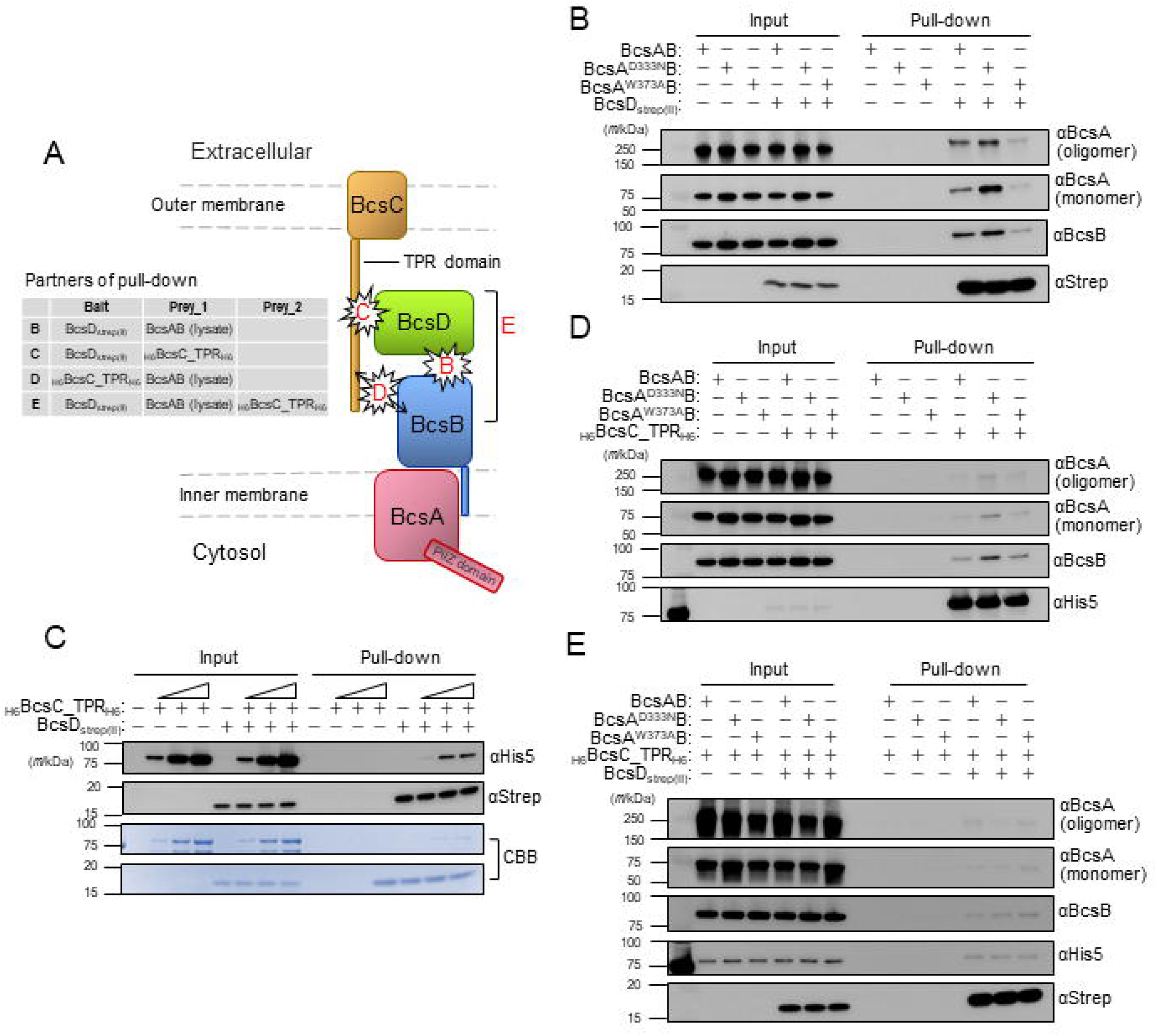
Pull-down assay with the Bcs subunits. **A** Schematic diagram showing the partners (bait and prey proteins) in a pull-down assay for each subunit of BcsABCD. Alphabets on the cartoon indicate panels B–D (the result of pull-down experiments), by which the subunit-subunit interaction was surveyed. **B** Western blot images of the pull-down assay using BcsD_StrepII_ (bait) with crude BcsAB (prey). The pull-down fraction was analyzed by western blot using anti-BcsA, anti-BcsB, and anti-Streptag antibodies. **C** Western blot images of the pull-down assay using BcsD_StrepII_ (bait) with purified _H6_BcsC_TPR_H6_ (prey). The pull-down fraction was analyzed by CBB staining and western blotting using anti-His5 and anti-Streptag antibodies. **D** Western blot images of the pull-down assay using _H6_BcsC_TPR_H6_ (bait) with crude BcsAB (prey). The pull-down fraction was analyzed by western blot using anti-BcsA, anti-BcsB, and anti-His5 antibodies. **E** Western blot images of the pull-down assay using BcsD_StrepII_ (bait) with crude BcsAB (prey_1) and purified _H6_BcsC_TPR_H6_ (prey_2). The pull-down fraction was analyzed by western blot using anti-BcsA, anti-BcsB, anti-His5, and anti-Streptag antibodies. Input: sample immediately after mixing; pull-down: elution fraction after pull-down. All the detailed methods are described in the “Experimental Procedures” section of the text.

First, the interaction between BcsAB and BcsD, which has been demonstrated above, was tested by pull-down with a bait of Streptag(II)-fused BcsD (BcsD_StrepII_) and a prey of crude BcsAB without tags (DDM-solubilized cell membrane of *E. coli* expressing BcsAB). As expected, BcsAB bound to BcsD_StrepII_ (Fig. 6B). The pull-down assay yielded a result consistent with our purification experiment of the BcsABD-complex and supported our hypothesis for the interaction between BcsAB and BcsD.

Given the successful pull-down assay with BcsAB and BcsD_StrepII_, we next performed a pull-down assay to survey the BcsC-involved interaction in the complex (see Fig. 6A). For this purpose, TPR tetratricopeptide-repeat (TPR) domain of the BcsC protein (BcsC_TPR) was used based on two assumptions: (i) the role of the TPR domain is protein-protein interaction (26), and (ii) the full-length BcsC protein is unstable, as shown in our purification. Pull-down by BcsD_StrepII_ trapped _H6_BcsC_TPR_H6_, which has hexa-histidine tags at both the N-and C-termini of BcsC_TPR, in a dose-dependent manner (Fig. 6C), clearly showing that these two proteins interact with each other. Furthermore, pull-down by the purified _H6_BcsC_TPR_H6_ (bait) with the crude BcsAB (prey) also supported their interaction with each other (Fig. 6D), although it appeared less striking than the case with _H6_BcsC_TPR_H6_ and BcsD_StrepII_ (Fig. 6C).

Finally, we performed a pull-down assay using all subunits in the complex: crude BcsAB, purified _H6_BcsC_TPR_H6_, and purified BcsD_StrepII_ (Fig. 6E). Streptactin beads trapped BcsA, BcsB, and _H6_BcsC_TPR_H6_ together with BcsD_StrepII_, although BcsA was only faintly detected by WB. It is then summarized that the BcsD protein can simultaneously interact with BcsA, BcsB, and the TPR region of BcsC.

## Discussion

Bacterial cellulose synthase is a heterosubunit complex. Primary evidence for this hypothesis is the existence of gene clusters or operons encoding cellulose synthase-related genes (3,6,7,11). However, this hypothetical model has not been tested intensively, owing to the lack of studies on the protein. Two recent reports have addressed this question. Sunagawa et al. demonstrated an interaction between BcsD and BcsH (Ccp) proteins using a pull-down assay (16). Another prominent study was the X-ray crystallographic analysis of the BcsAB protein complex by Morgan et al., which clarified that the BcsA and BcsB proteins closely interact with each other to polymerize and translocate the glucan chain from the cytosol to the extracellular side (1), as confirmed also in *E.coli* (27). In this study, we analyzed the type I Bcs complex and demonstrated that the BcsD protein interacted with the BcsAB complex (Fig. 3, 4, 5, and Fig. 6B) and the TPR domain of BcsC (Fig. 6C). Furthermore, it was shown that BcsC_TPR interacted with the BcsAB complex, although the result was not as clear as the interaction between BcsC_TPR and BcsD (Fig. 6D). Finally, a pull-down assay using BcsD_StrepII_ with a mixture of crude BcsAB and purified _H6_BcsC_TPR _H6_ indicated that BcsD binds to both BcsAB and BcsC_TPR simultaneously, suggesting that these four subunits interact with each other (Fig. 6E). Our study, together with previous studies, experimentally demonstrated that the subunits of BcsA, BcsB, BcsC, BcsD, and BcsH are included in the type I Bcs-complex.

Interestingly, this study showed that the interaction of BcsD with BcsAB has two modes: direct and indirect interactions through the polymerized glucan chain (Fig. 4). Direct interaction between the BcsAB complex and BcsD protein was observed for both wild type and catalytically deficient mutant cellulose synthase, whereas the interaction via cellulose was observed only for the wild type (Fig. 5). We then propose a two-state model for the type I Bcs complex: basal and active states, which are represented by the direct interaction (Fig. 7, left) and indirect interaction modes (Fig. 7, right), respectively.

**Figure 7.**
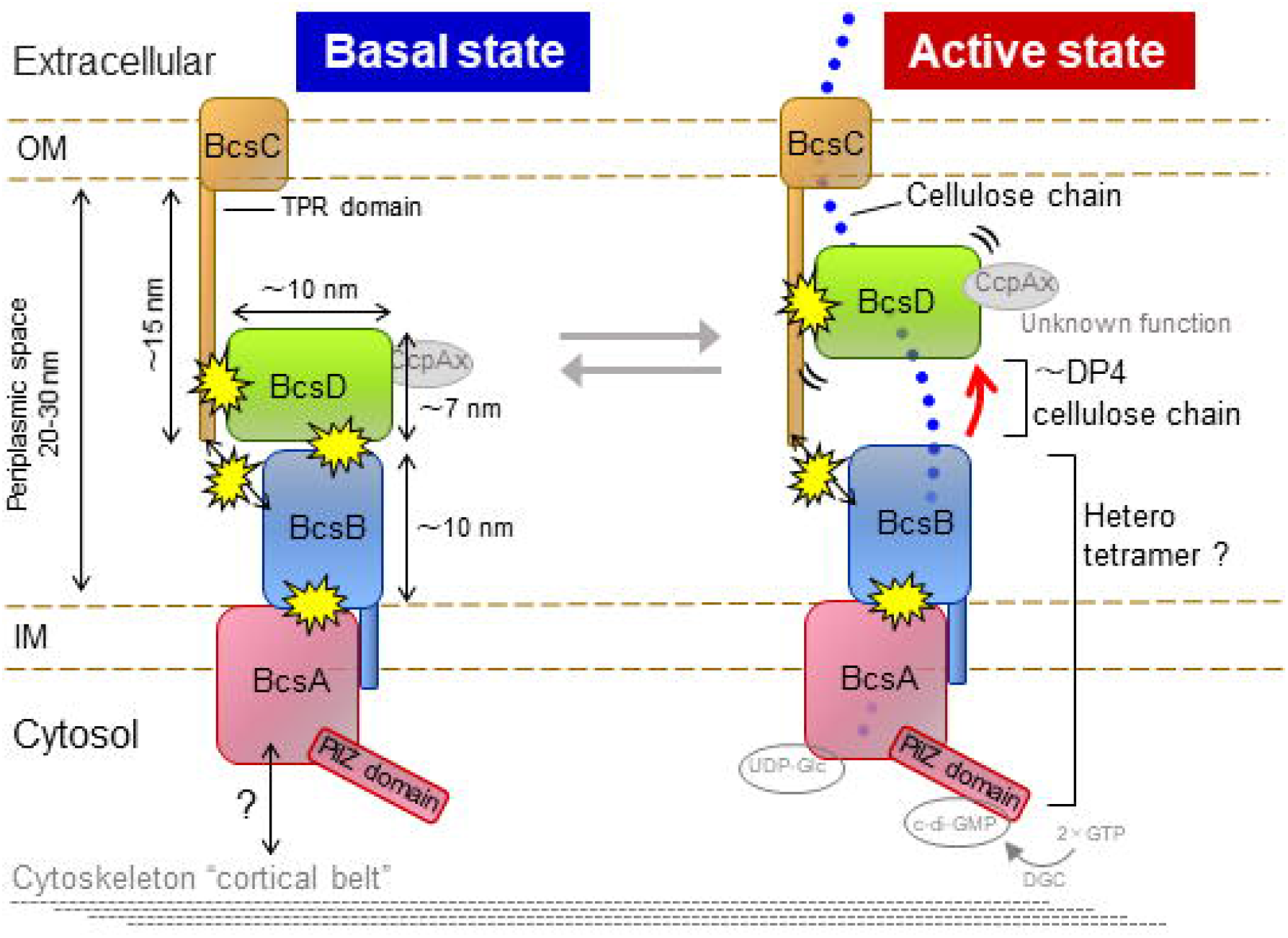
The hypothetical two-state model for the type I Bcs-complex. Hypothetical two-state model for the Bcs-complex: basal state (left) and active state (right). In this model, BcsD plays a key role in the periplasmic space to control the state of the complex separately from the state of the BcsA enzyme (1, 2). In the active state (right), BcsD draws the glucan chains in its octameric ring and brings those chains to the channel pore of BcsC in the outer membrane like a “button whirligig’. The TPR domain in BcsC may escort the BcsD protein to correctly transfer the glucan chains in BcsD to the BcsC channel. In the basal state (left), BcsD docks with the BcsB complex given that BcsB is located in the periplasmic space (1) as well as BcsD (12, 20). In the model, BcsD in the basal state also docks with BcsC without interfering with BcsB and is ready to draw the glucan chains in it once BcsAB starts synthesizing and extruding out the glucan chain. Thus, BcsD in this state may be a standby state for effectively transferring glucan chains into the extracellular space through the BcsC channel once BcsA is activated. Subunits that were not assessed in this study are indicated in gray.

BcsD protein in this basal state is statically fixed on the BcsAB complex for lying in wait for a glucan chain synthesized by the BcsAB complex. Once cellulose synthesis is initiated, the synthesized glucan chain is guided into the octameric ring of BcsD to crystallize cellulose molecules, as proposed previously (7,15,16,20,28), which is a shift from the resting state to the active state of the complex. In the active state (Fig. 7, right), the BcsD protein is detached from the BcsAB complex but tethered to BcsAB by the glucan chain extruded from BcsAB. Considering the substrate specificity of BsCel5A (Fig. S1), BcsD can be physically separated from BcsB by a glucan chain with a length of four or more glucose residues (Fig. 7).

In this two-state model, the active state of the Bcs complex allows BcsD to be launched toward the outer membrane, while BcsD in the basal state may be fixed statically on BcsB. The TPR domain of the BcsC protein may control diffusible BcsD in the active state in the periplasmic space by gentle but significant interactions. Given that the C-terminal domain of BcsC is an outer membrane β-barrel channel (Fig. S4) (24), the TPR domain of BcsC may escort BcsD carrying glucan chains toward the outer membrane channel in BcsC to spin the glucan chains to the extracellular side. The following questions about this model will be whether (i) BcsD protein could dock to the outer membrane channel part of the BcsC protein and (ii) the interaction between BcsC and BcsD could be state-dependent. These questions will be addressed in future studies to enhance our understanding of the Bcs-complex function.

Our study demonstrated the intermolecular interactions of subunits in the BcsABCD complex to update our knowledge of the quaternary structure of the type I Bcs complex. Experiments in this study were performed using recombinant proteins heterogeneously expressed in *E. coli*; thus, further studies will be necessary to test the hypothesis proposed in this study, the two modes for BcsAB-BcsD interaction: the basal and the active state. Nonetheless, the data obtained in this study were intensively tested using mutagenesis experiments of proteins, and they were consistent with each other. We believe that our hypothesis in this study is sufficiently reliable, at least for starting further discussion about the mechanism of Bcs with regard to protein function. In this model, BcsD is not statically fixed in the Bcs-complex but dynamically interacts with the other subunits in the complex (Fig. 7). Such a dynamic behavior may be involved in the crystallization function of cellulose molecules, as well as twisting in bacterial cellulose microfibrils, as reported previously (16). The putative state control with BcsD, as shown in Fig. 7, is the regulation of the quaternary structure of bacterial cellulose synthase. Next, we need to further study the original *Acetobacter* cells and observe the whole Bcs-complex at higher resolution with cryo-EM under controlled two-state conditions.

### Experimental procedures

#### Chemicals

All chemicals used in this study were purchased from FUJIFILM Wako Pure Chemicals Co. Ltd. (Osaka, Japan), Nacalai Tesque Co. Ltd. (Kyoto, Japan), and Sigma-Aldrich Inc. (MO, USA), unless otherwise indicated.

#### Cloning

Bcs genes (*bcsA*, *bcsB*, *bcsC*, and *bcsD*) were prepared from genomic DNA extracted from cultured cells of *Komagataeibacter sucrofermentans* JCM9730, which was provided by the RIKEN Bio Resource Center (Tsukuba, Japan) and known by the strain name BPR2001 (29, 30). The endo-β1,4-glucanase gene from *Bacillus subtilis* strain 168 (*bscel5A*) (23) was prepared from the genomic DNA provided by the American Type Culture Collection (ATCC23857D). The dgc gene (*dgc*) was identical to that used in our previous study (18): tDGC and DGC from *Thermotoga maritima* MSB8 with a point mutation of R158A and truncation of the N-terminal 81 amino acid residues from V1 to V81 (31). The gene for green fluorescent protein UV (GFPuv) (32) was prepared from the pGFPuv plasmid purchased from Takara Bio, Co. Ltd. (Shiga, Japan).

The cloning of these genes was performed by PCR using PrimeStarGXL (Takara Bio Co. Ltd.). Site-directed mutagenesis was performed by QuikChange II (Agilent Inc., CA, USA) or the equivalent PCR-based method. The sequence of the PCR-amplified DNA was checked by the dye-terminator method in a subcloning vector (pGEM-T easy (Promega Inc., WI, USA) or pTAC-2 (BioDynamics Laboratory Co. Ltd., Tokyo, Japan)), or final expression vector, which was performed at either of the Uji DNA-sequencing core (Kyoto University Research Administration Office) with an ABI Prism 310 Genetic Analyzer (Applied Biosystems Inc., MA, USA), or Eurofins Genomics Inc. (Tokyo, Japan).

#### Construction of the expression vectors

The protein expression vectors used in this study are summarized in Table 1. Expression vectors for BcsAB (BcsAB, BcsA^D333N^B, and BcsA^W373A^B) were identical to those used in previous studies (18, 21). The *bcsAB* gene in the operon form was inserted between the EcoRI and KpnI sites in pQE-80L (Qiagen, Hilden, Germany), and the upstream of the start codon was the original ribosomal binding site of *K. sucrofermentans* JCM9730.

**Table 1.**
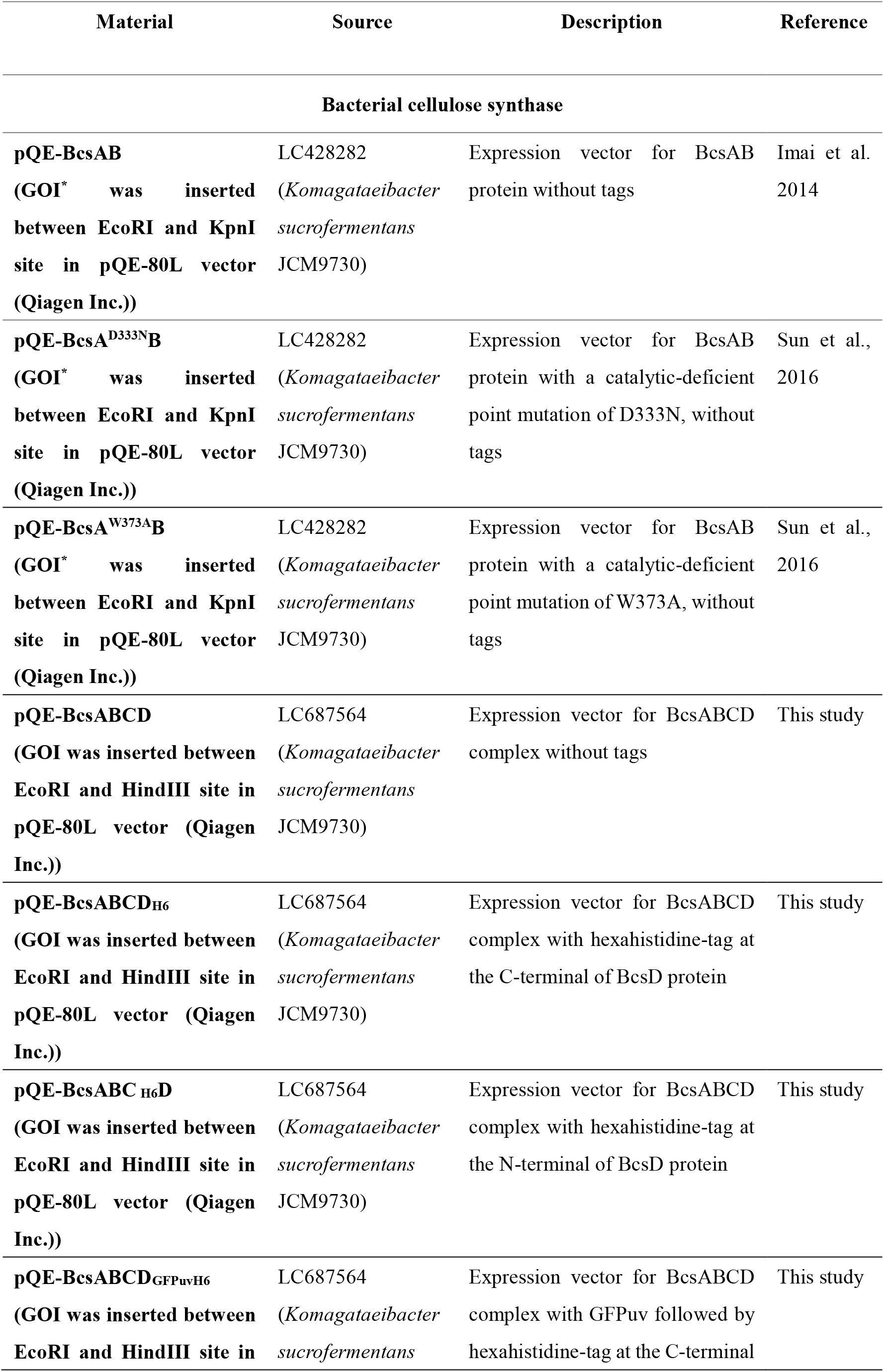
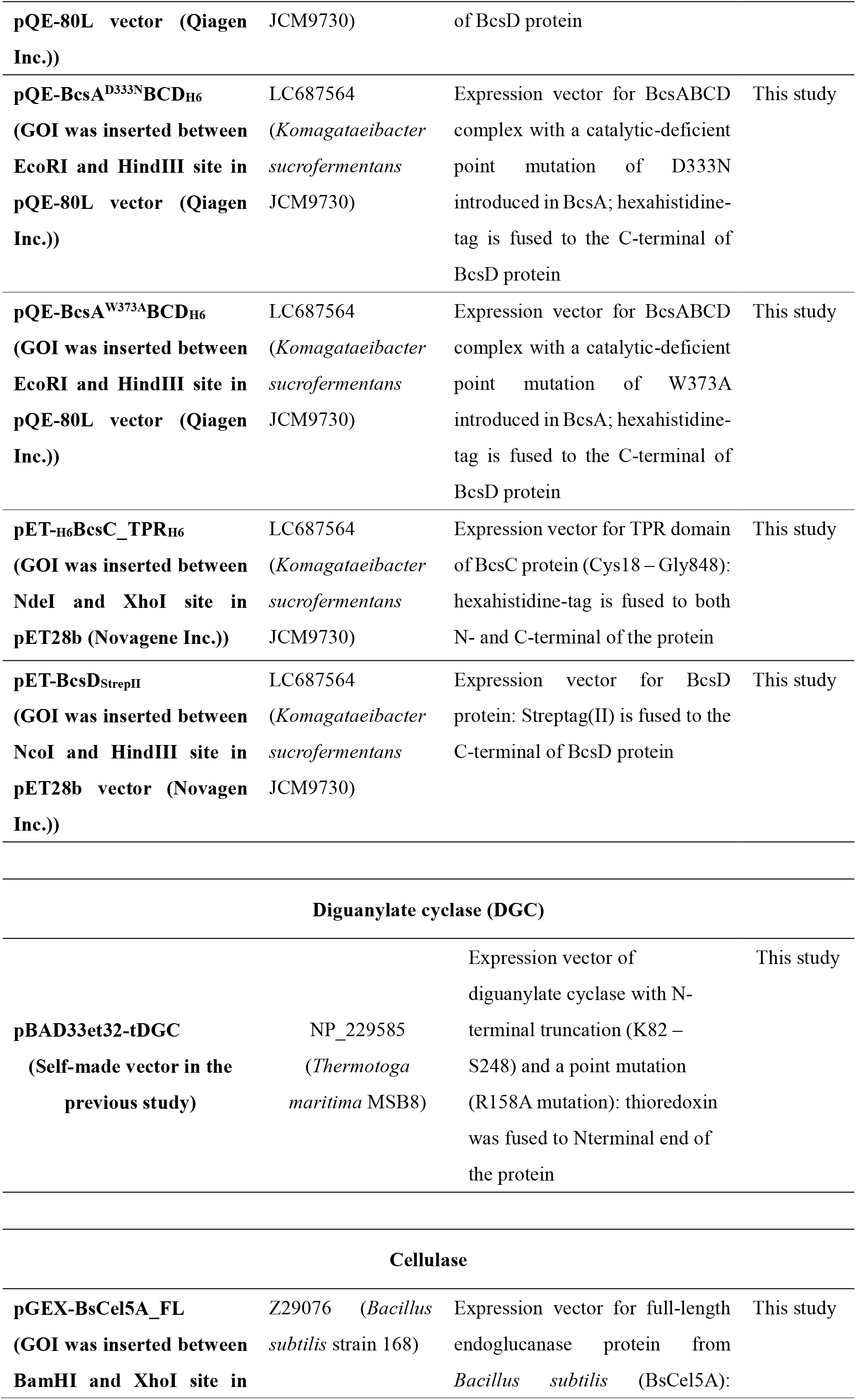
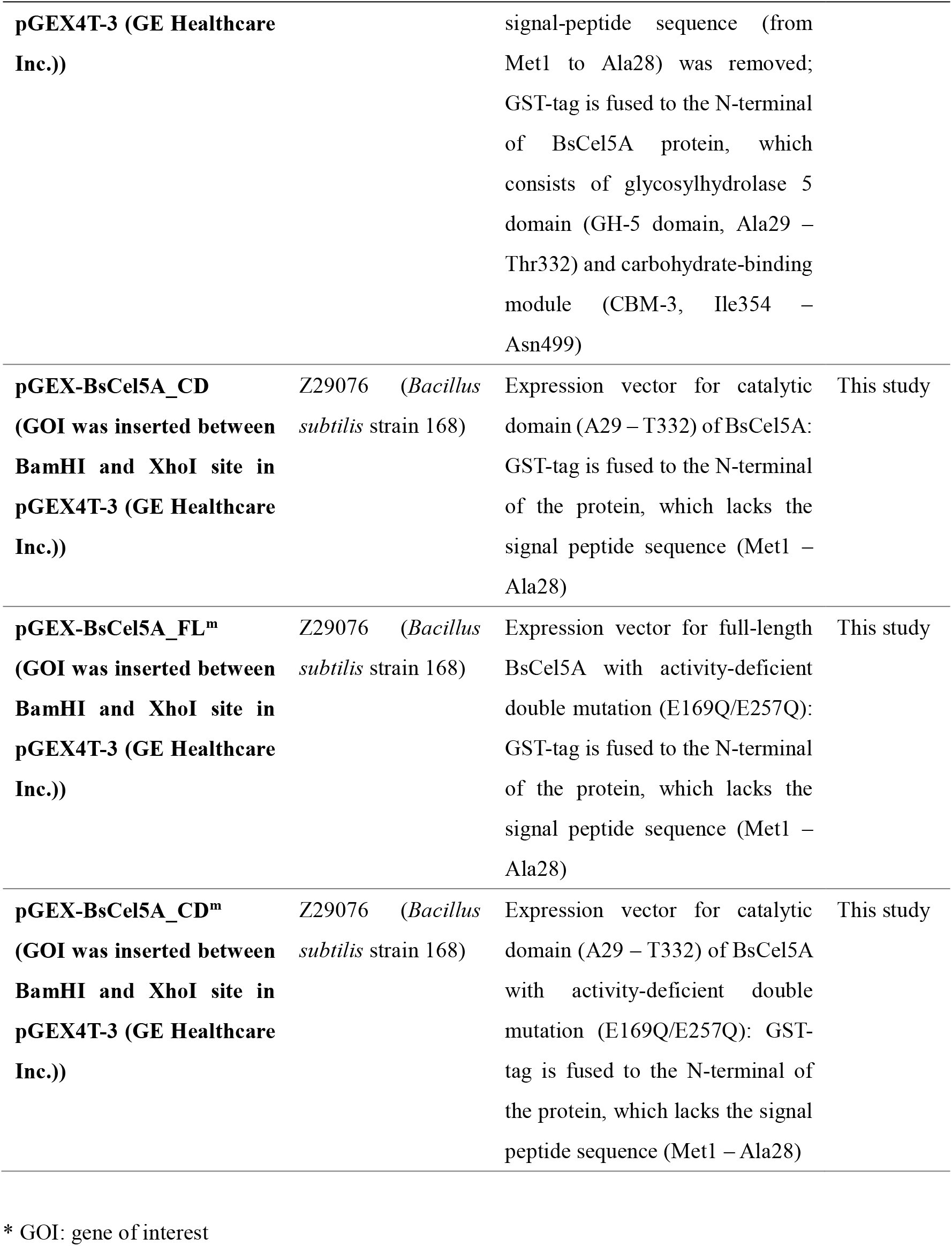
Experimental resources to express the proteins in this study.

Expression vectors for BcsABCD (BcsABCD, BcsABC_H6_D, BcsABCD_H6_, BcsA^D333N^BCD_H6_, BcsA^W373A^BCD_H6_, and BcsABCD_GFPuvH6_) proteins were constructed from the vectors for BcsAB by inserting the *bcsABCD* gene in the operon form between the EcoRI and HindIII sites in pQE-80L. The ribosomal binding site in these BcsABCD expression vectors is exactly the same as that in the expression vectors of BcsAB, one of *K. sucrofermentans* JCM9730. DNA encoding the tag (hexa-histidine or GFPuv) was added to *bcsD* during the PCR-amplification amplification.

The expression vectors for the TPR region of BcsC (BcsC_TPR: C18-G848) and BcsD were prepared for the pull-down assay. DNA encoding hexa-histidine tag was added to the 3′-end of the gene for BcsC_TPR during PCR amplification and inserted into the NdeI and XhoI sites in pET-28b (Merck Corp.., Darmstadt, Germany) to express BcsC_TPR protein with dual hexa-histidine tag at both N-and C-terminal – _H6_BcsC_TPR_H6_. DNA encoding Streptag (II) was added to the 3′-end of the gene for BcsD and inserted between the NcoI and HindIII sites in pET28b to express BcsD with C-terminal Streptag(II)–BcsD_StrepII_.

The gene for tDGC was inserted between the NdeI and HindIII sites in pBAD33et32, a customized plasmid DNA in this study, to express the thioredoxin-fused tDGC protein to count the cellulose-synthesizing activity of the recombinant BcsAB and BcsABCD complex in our previous studies (18, 21). The detailed procedure for preparing pBAD33et32 is described in Supplementary Information. Thioredoxin gene (*trx*) in pBAD33et32 was copied from pET32a (Merck Corp.) to replace the His-patch thioredoxin used in our previous study (18), which was derived from pBAD202-TOPO (Invitrogen Inc., MA, USA).

For the bacterial endoglucanase gene bscel5A, the putative signal peptide sequence (M1–A28) was removed from the full-length construct. Full-length (A29–N499) and catalytic domain (A29–T332) genes were inserted between BamHI and XhoI sites in pGEX4T-3 (Cytiva Inc., MA, USA) to express the N-terminal GST-fused endoglucanase BsCel5A.

#### Deletion of bcsA gene from Escherichia coli BL21

In this study, we used genetically modified *E. coli* BL21 to express Bcs proteins, which lack the whole *bcsA* gene (Fig. 1A). This gene manipulation was performed using the Red/ET recombination system (Gene Bridges GmbH, Heidelberg, Germany), according to the manufacturer’s specifications. The detailed procedure is described in Supplementary Information.

Briefly, *E. coli* BL21 strain (purchased from New England Labs Inc., MA, USA) was transformed with the pRedET plasmid carrying the genes for DNA-recombination proteins and selected by tetracycline resistance. The second transformation was followed by introducing linear DNA of neomycin-resistant gene flanked by the homology arm sequences corresponding to the upstream and downstream of *bcsA* gene (CP010816.1); the recombinant proteins were expressed with arabinose-induction for disrupting *bcsA* gene, and subsequently the pRedET plasmid was expelled out of the cell by culturing at 37°C. Finally, the *bcsA-disrupted* BL21 transformant was selected for kanamycin resistance and tetracycline sensitivity. Gene modification was confirmed by direct PCR (Fig. 1B). We named this genetically modified *E. coli* BL21 strain as BL21-GM. Chemically competent cells of BL21-GM were then prepared by treating the cultured cells in a rubidium-based buffer according to the protocol on the website of the Center for Molecular Biology and Genetics at Mie University (https://www.gene.mie-u.ac.jp/Protocol/Original/Transform-Comp-Cell-Freeze.html, in Japanese) (33).

#### *In vivo* cellulose synthase activity

The endogenous cellulose-synthesizing activity of *E. coli* and cellulose-synthesizing activity of the heterogeneously expressed Bcs-complex in *E. coli* was estimated by expressing DGC protein to activate bacterial cellulose synthase in the cell by the produced c-di-GMP (18). In this study, the pBAD33et32-tDGC plasmid was used to express the DGC. The obtained *E. coli* transformants were pre-cultured in LB medium including the proper antibiotics at 37°C overnight, and then 0.1 mL of the pre-culture was inoculated into 10 mL of ZYP-5052 medium with 50 μg/mL chloramphenicol (100 μg/mL ampicillin when expressing Bcs-complex) and 0.001% L-arabinose. Cells were grown at 20°C in an orbital shaker at 180 rpm for 24 h and the cultured cells were collected by centrifugation at 2,000 × *g* for 20 min. The amount of cellulose in the pellet was quantified using the anthrone-sulfuric acid method (34) as described below.

Two milliliters of AN-reagent (a mixture of glacial acetic acid and nitric acid with volumetric ratio of 8:1) were added to the pellet and then heated at 105°C for 30 min. The remaining cellulose pellets were washed with 10 mL of water once by centrifugation at 2,000 × *g* for 10 min, and then 1 mL of 72% (v/v) H_2_SO_4_ was added to the pellets and incubated for 1 h at room temperature. The solution (0.1 mL) was mixed with water (0.4 mL), and then, 1 mL of fresh anthrone reagent (0.2% w/v anthrone in concentrated H_2_SO_4_) was added and mixed. The solution was heated to 90°C for 15 min and allowed to stand until it reached room temperature (approximately 15 min). The absorbance of the solution was measured at both 625 and 700 nm, and the absorbance at 625 nm, from which the absorbance at 700 nm was subtracted, was used for quantification for better reproducibility. Calibration was performed with phosphoric acid-swollen cellulose (PASC) prepared using a previously reported protocol (35).

#### Purification and characterization of BsCel5A

Expression of GST-fused BsCel5A endoglucanase was performed in *E. coli* BL21 (not GM strain): all the four constructs ((BsCel5A_FL, BsCel5A_CD, BsCel5A_FL^m^, and BsCel5A_CD^m^) were expressed and purified in the same manner.

The transformant of BL21 with pGEX-BsCel5A were inoculated in 10 mL LB containing 100 μg/mL ampicillin in a 100 mL flask and grown at 37°C with orbital shaking at 220 rpm overnight. The cells were harvested by centrifugation at 3,000 × *g* for 20 min, suspended in 1 L of LB containing 100 μg/mL ampicillin in a 5 L baffle flask, and cultured at 37°C until OD_600_ reached 0.6. The culture medium was cooled in ice water for 10 min, and then IPTG was added to 0.1 mM and incubated at 20°C for 20 h at 160 rpm with shaking.

The cells were then collected at 3,000 × *g* for 20 min, washed with 40 mL of 1 × phosphate-buffered saline (PBS), suspended in 40 mL of 1 × PBS containing 1 mM phenylmethylsulfonyl fluoride (PMSF), and sonicated on ice for ∼5 min (exposure: 10 s, rest: 50 s; until slightly translucent). The cell lysate clarified by 20,000 × *g* for 30 min at 2°C was applied to 5 mL of Glutathione Sepharose 4B (Cytiva Inc,), washed with 20 mL of 1 × PBS, and eluted with 5 c.v. of 0.1 M Tris-HCl pH 8.0, 0.5 M NaCl, 20 mM reduced-glutathione. The eluted fraction was buffer exchanged with 1 × PBS using Vivaspin 6 (MWCO 10 kDa; Sartorius AG, Germany).

To remove the GST tag fused to the N-terminus, 100 units of thrombin (Merck Corp.) was added to 10 mg of _GST_BsCel5A and incubated at 20°C for two days. Thrombin was then removed by adding 1 mL of Benzamidine Sepharose 6 B (Cytiva Inc.), and the cleaved GST was removed by adding 5 mL of Glutathione Sepharose 4 B. A solution of PMSF was added to the obtained BsCel5A protein at a final concentration of 1 mM to inactivate a small amount of residual thrombin.

The recombinant BsCel5A proteins purified were characterized with regard to the following points: (i) binding activity to cellulose, (ii) glycoside hydrolase activity, and (iii) substrate specificity. The detailed experimental procedure is described in Supplementary Information.

#### Purification of BcsABCD complex

All the protein purification experiments in this study were performed at low temperature (∼4°C). The composition of the protease inhibitor mix (PI) used for protein purification was as follows: 0.1 mM PMSF, 1 mM benzamidine-HCl, 5 µM bestatin, 10 µM leupeptin, 1 μM pepstatin A.

The transformant of *E. coli* BL21-GM with pQE-BcsABCD_H6_ was inoculated in 10 mL of LB containing 100 μg/mL ampicillin in a 100 mL flask and grown at 37°C with shaking at 220 rpm overnight. The cells were harvested by centrifugation at 3,000 × *g* for 20 min, resuspended in 1 L of ZYP-5052 containing 100 μg/mL ampicillin and 0.0005% (v/v) antifoam SI (Fujifim Wako Chemicals Co. Ltd., Japan) in a 5 L baffle flask, and incubated at 20°C for 22 h with an orbital shaker at 100 rpm. The cultured cells were collected at 3,000 × *g* for 20 min and suspended in 50 mL of buffer A (20 mM sodium phosphate buffer (pH 7.5), 300 mM NaCl, 8% glycerol, 5 mM MgCl_2_, 5 mM cellobiose) with PI, prior to cell lysis by two passes of high-pressure homogenizer (French Press, Thermo Fisher Scientific Inc., MA, USA) at a pressure of 15,000 psi. The lysate clarified by centrifugation at 5,000 × *g* for 10 min and ultracentrifugation at 150,000 × *g* for 30 min at 2°C. The obtained cell membranes were resuspended in buffer A with PI and NaCl concentrations increased to 1 M and were ultracentrifuged again under the same conditions. Finally, the cell membrane was suspended in buffer A with PI to approximately 10 mg/mL (estimated by the DC protein assay, Bio-Rad Inc., CA, USA). The collected cell membrane was stored at −80°C until use in experiments.

Ten milliliters of the cell membrane suspension were mixed with *n*-dodecyl-β-D-maltopyranoside (DDM) at a final concentration of 0.5% (w/v) and shaken gently for 1 h at 4°C for solubilizing Bcs-complex. Supernatant of the following ultracentrifugation at 150,000 × *g* for 30 min at 2°C was collected as solubilized fraction, and imidazole stock solution (2 M, pH 8.0) was added to achieve a final concentration of 10 mM, prior to loading to 0.4 mL a His-Select Cobalt Affinity Gel (Sigma-Aldrich Inc.), which was equilibrated with 4 mL of wash buffer (20 mM imidazole and 0.025% (w/v) DDM in buffer A) in advance. After loading the solubilized protein onto the affinity resin in a plastic open column, the resin in the column was washed with 3 mL of wash buffer and finally eluted with elution buffer (200 mM imidazole and 0.025% (w/v) DDM in buffer A).

#### Cellulase treatment of Bcs-complex

Cellulase treatment of BcsABCD_H6_ was performed using a cobalt affinity gel. The same procedure was followed for the binding step of BcsABCD_H6_ on the affinity gel. After all the solubilized fraction passed through the column, 0.8 mL of 1.5 μM BsCel5A solution in wash buffer (20 mM imidazole and 0.025% (w/v) DDM in buffer A) was loaded to the column. The column exit was plugged when most of the cellulase solution passed, and then the gel was incubated at 10°C for 3 h with the remaining cellulase solution. After unplugging, the affinity gel in the column was washed with 2 mL of wash buffer and eluted with elution buffer (200 mM imidazole and 0.025% (w/v) DDM in buffer A).

#### Protein preparation for pull-down assay: BcsC_TPR, BcsD, and crude BcsAB

The pull-down assay in this study used purified BcsC_TPR, purified BcsD, and crude BcsAB proteins, each of which was prepared as described below.

The transformant of *E. coli* BL21(DE3) with pET-_H6_BcsC_TPR_H6_ was inoculated in 10 mL of LB containing 50 μg/mL kanamycin in a 100 mL flask and grown at 37°C with shaking at 220 rpm overnight. The cells were harvested by centrifugation at 3,000 × *g* for 20 min, and resuspended in 1 L of ZYP-5052 (36) containing 200 μg/mL kanamycin and 0.0005%(v/v) antifoam SI in a 5 L baffle flask, and incubated at 20°C for 24 h in a shaking incubator at 100 rpm. The cultured cells were collected at 3,000 × *g* for 20 min, and washed with 40 mL of 1 × PBS prior to resuspension in 100 mL of 10 mM imidazole in buffer B (20 mM sodium phosphate buffer (pH 7.5), 500 mM NaCl, 8% glycerol) with PI, and sonicated on ice for ∼5 min (exposure: 10 s, rest: 50 s; until slightly translucent). The cell lysate was clarified by centrifugation at 40,000 × *g* for 30 min at 2°C and applied to 4 mL a His-Select Cobalt Affinity Gel (Sigma-Aldrich Inc.), followed by washing with 10 mM imidazole in buffer B, and eluted 250 mM imidazole in buffer B. Finally, the buffer of eluted fraction including _H6_BcsC_TPR_H6_ protein was replaced with buffer B using PD-10 desalting column (Cytiva Inc.).

The BcsD protein was expressed in *E. coli* BL21(DE3) with pET-BcsD_StrepII_. The transformant was inoculated in 10 mL of LB containing 50 μg/mL kanamycin in a 100 mL flask and grown at 37°C with shaking at 220 rpm overnight. The cells were harvested by centrifugation at 3,000 × *g*, 20 min, resuspended in 1 L of LB containing 50 μg/mL kanamycin in a 5 L baffle flask and cultured at 37°C until OD_600_ was ∼0.6. The culture medium was cooled in ice water for 30 min, and IPTG (0.5 mM) was added and incubated at 25°C for 20 h at 120 rpm. Cells were collected at 3,000 × *g* for 20 min, suspended in 30 mL of buffer C (20 mM Tris-HCl (pH 7.5), 150 mM NaCl, and 1 mM EDTA) with PI, and sonicated on ice for ∼10 min (exposure: 10 s, rest: 50 s; until slightly translucent). The cell lysate was centrifuged at 20,000 × *g* for 30 min at 2°C, applied to 2 mL of Streptactin Sepharose HP (Cytiva Inc.), washed with 20 mL of buffer C, and eluted with 4 mM *d*-desthiobiotin in buffer C. Finally, buffer of eluted fraction was replaced by buffer C using Amicon Ultra-4 MWCO 50 kDa (Merck Corp.).

The crude protein of BcsAB was prepared using the transformant *E. coli* BL21-GM with pQE carrying the *bcsAB* gene. The procedure to isolate the cell membrane, from culture to ultracentrifugation, was same as in the case of BcsABCD_H6_ as described previously, except for the buffer to suspend the cells and cell membrane: buffer D (20 mM sodium phosphate buffer (pH 7.5), 100 mM NaCl, 8% glycerol, 5 mM MgCl_2_, and 5 mM cellobiose) with PI was used. The obtained cell membrane was suspended in buffer D with PI at approximately 10 mg/mL (estimated by the DC protein assay, Bio-Rad Inc.). Finally, crude BcsAB protein was prepared by solubilizing 7 mL of the cell membrane fraction with 0.5% (w/v) DDM for 1 h and centrifugation at 150,000 ×*g* for 30 min at 2℃.

#### Pull-down assays

For the pull-down by BcsD_StrepII_ with the crude BcsAB, purified BcsD_StrepII_ (0.2 mg) was added to 7 mL of crude BcsAB, incubated for 3 h at 4 ℃, and then loaded to 0.1 mL of Streptactin Sepharose HP. The resin was washed with 1 mL of 0.025% (w/v) DDM in buffer D, and finally the bound protein was eluted with 4 mM desthiobiotin in buffer D with 0.025% (w/v) DDM.

For the pull-down assay, BcsD_StrepII_ with _H6_BcsC_TPR_H6_, BcsD_StrepII_ (0.1 mg) were incubated with _H6_BcsC_TPR_H6_ (0, 0.1, 0.3, 0.5 mg) in 3 mL of buffer D (–/–) (buffer D without MgCl_2_ and cellobiose) for 3 h at 4 ℃, and then loaded to 0.1 mL of Streptactin sepharose HP. After washing with buffer D (–/–), the bound protein was eluted with 1 mL 4 mM *d*-desthiobiotin in buffer D (–/–).

For the pull-down assay by _H6_BcsC_TPR_H6_ with BcsAB, purified _H6_BcsC_TPR_H6_ (0.2 mg) was mixed with crude BcsAB (7 mL), and further NaCl stock solution was added to set 500 mM NaCl in the final solution. After incubation for 1 h at 4 ℃, the sample was loaded to 0.2 mL a His-Select Cobalt Affinity Gel and washed with 2 mL of 20 mM imidazole in buffer B (+/+) (buffer B with 5 mM MgCl_2_ and 5 mM cellobiose) with 0.025% (w/v) DDM. The bound protein was eluted with 200 mM imidazole in buffer B containing 0.025% (w/v) DDM.

Finally, we performed a pull-down assay using BcsD_StrepII_, purified BcsC_TPR; crude BcsAB. BcsD_StrepII_ (0.2 mg) and _H6_BcsC_TPR_H6_ (0.2 mg) were mixed with crude BcsAB (7 mL of DDM extract); NaCl stock solution (5 M) was added to a final concentration of 500 mM NaCl. After incubation for 3 h at 4 ℃, the sample solution was loaded to 0.1 mL of Streptactin Sepharose HP. After washing with 1 mL of buffer B (+/+) containing 0.025% (w/v) DDM, the bound protein was eluted with 4 mM *d*-desthiobiotin in buffer B (+/+) containing 0.025% (w/v) DDM.

Prey protein without bait protein was used as a negative control for the pull-down assay using the same procedures. The eluted fraction was analyzed by SDS-PAGE, followed by WB as described later.

#### Western blotting (WB) analysis

Rabbit polyclonal antibodies against BcsA, BcsB, BcsC, and BcsD proteins were generated with the synthetic peptide corresponding to a part of the C-terminus of BcsA (18,20,21,37), the C-terminus of BcsB (this study), the C-terminus of BcsC (37), and the loop between the β3 and β4-strand (20, 37). The dilution rate was 10,000-fold for anti-BcsA, anti-BcsB, and anti-BcsC, whereas anti-BcsD was diluted 20,000-fold. Each of the antibodies against the peptide tag were diluted as follows: anti-Strep tag II (15 ng/mL; ab180957, Abcam Corp., Cambridge, UK), anti-His5 (0.1 μg/mL; 34660, Qiagen N.V., Germany), and anti-Thioredoxin (1 μg/mL; ab139677, Abcam Corp.).

Horseradish peroxidase (HRP)-conjugated anti-rabbit IgG (W401B; Promega Inc.) and HRP-conjugated anti-mouse IgG (7076S; Cell Signaling Technology Inc., MA, USA) were diluted 20,000-fold and used as the secondary antibodies. ECL-Prime (Cytiva Inc.) was used for chemical luminescence with HRP.

For the WB with anti-BcsD, anti-Strep, anti-His, and anti-thioredoxin, the samples for SDS-PAGE were heat-treated at 100°C for 3 min after mixing with the sample buffer, while the samples to visualize by the other antibodies were incubated in the sample buffer for several minutes at room temperature. The samples were analyzed by SDS-PAGE with a precast polyacrylamide gel (SuperSep Ace 5-20%, Fujifim Wako Chemicals Co. Ltd.), and blotting from the gel to PVDF membrane (Immobilon-P, 0.45 μm-ϕ, Cytiva Inc.) was done by a constant current mode at 150 mA/10 cm^2^ for 30 min. Blocking of the membrane was performed with 0.3% skim milk and 0.02% NaN_3_ in tris-buffered saline (TBS) containing 0.05% Tween-20 (TBS-T), for the WB with anti-Bcs proteins, while blocking was performed with 2.5% BSA, 0.1% gelatin, 0.02% Tween-20, 0.04% NaN_3_ in PBS at 4°C overnight or at room temperature for 0.5–1 h, for either of anti-Strep-tag II, anti-His5, or anti-thioredoxin. The membranes were briefly rinsed with TBS-T and then subjected to the primary antibody reaction in blocking buffer for 1–2 h at room temperature. After removing the antibody solution, the membrane was briefly rinsed and washed with TBS-T thrice for 5 min, followed by a secondary antibody reaction in TBS-T for 30–45 min at room temperature. The membrane was briefly rinsed and washed with TBS-T three times for 15 min each time. The protein bands were then visualized by casting the ECL prime solution on the blotted PVDF membrane (0.2 mL/9 cm2 membrane), and digitally recorded with AE-9300H EZ-Capture MG (ATTO Co. Ltd., Japan).

Quantitative analysis was performed using ImageJ (version 1.53a) to quantify the band signal intensity, calculated as relative values to the negative control, and the relative signal intensities of BcsA and BcsB were normalized to the signal intensity of BcsD. For BcsA; two major bands were reproducibly observed near the 75-kDa and 250-kDa marker (Fig. 2B, top row), as well as in our previous studies (18, 37), which are probably BcsA monomer and BcsA-including oligomer, respectively (the estimated molecular weight for the BcsA monomer is 84 kDa). Given these observations, the band intensity for the BcsA-containing oligomer was also considered in addition to the monomer band intensity in subsequent analyses.

#### Data availability

All data are contained within this article and available from the corresponding author on reasonable request.

## Supporting information

This article contains supporting information.

## Acknowledgments

The authors thank Dr. Kenji Tajima of Hokkaido University for his advice. The authors thank Professor Ryoichi Yamaji and Dr. Yasuyuki Kobayashi at Osaka Prefecture University for their advice regarding the pull-down assay.

## Author contributions

T. I., and M. Y. conceptualization; T. K., T. I., and M. Y. methodology; T. K. formal analysis, T. K., N. Y., and S. N. investigation; T. I. resources; T. K., and T. I. data curation; T. K., and T. I. writing-original draft; T. K., M. Y., and T. I. writing-review and editing; T. K., and T. I. visualization; T. I., and M. Y. supervision; T. K., M. Y., and T. I. project administration; T. I. funding acquisition.

## Funding and additional information

This work was supported by JPSP KAKENHI (grant numbers 15H04530 and 19H00950), Exploratory Research on Humanosphere Science (RISH, Kyoto University), and JST CREST (grant number: JPMJCR13B2).

## Abbreviations

The abbreviations used are:

Bcs: bacterial cellulose synthase
CD: catalytic domain
c.v.: column volume
c-di-GMP: cyclic-di-guanosine monophosphat
DDM: *n*-β-D-maltopyranoside
DGC: diguanylate cyclase
DP: degree of polymerization
FL: full length
GH: glycoside hydrolase
IMAC: immobilized metal affinity chromatography
PASC: phosphoric acid swollen cellulose
TBS: tris-buffered saline
TBS-T: tris-buffered saline with Tween-20
TPR: tetratricopeptide repeat

## References

1. Morgan, J. L. W., Strumillo, J., and Zimmer, J. (2013) Crystallographic snapshot of cellulose synthesis and membrane translocation. Nature 493, 181–186

2. Morgan, J. L. W., McNamara, J. T., and Zimmer, J. (2014) Mechanism of activation of bacterial cellulose synthase by cyclic di-GMP. Nat. Struct. Mol. Biol. 21, 489–496

3. Römling, U., and Galperin, M. Y. (2015) Bacterial cellulose biosynthesis: Diversity of operons, subunits, products, and functions. Trends Microbiol. 23, 545–557

4. Thongsomboon, W., Serra, D. O., Possling, A., Hadjineophytou, C., Hengge, R., and Cegelski, L. (2018) Phosphoethanolamine cellulose: A naturally produced chemically modified cellulose. Science 359, 334–338

5. Tajima, K., Imai, T., Yui, T., Yao, M., and Saxena, I. (2022) Cellulose-synthesizing machinery in bacteria. Cellulose 29, 2755–2777

6. Wong, H. C., Fear, A. L., Calhoon, R. D., Eichinger, G. H., Mayer, R., Amikam, D., Benziman, M., Gelfand, D. H., Meade, J. H., Emerick, A. W., Bruner, R., Ben-Bassat, A., and Tal, R. (1990) Genetic organization of the cellulose synthase operon in *Acetobacter xylinum*. Proc. Natl. Acad. Sci. U.S.A. 87, 8130–8134

7. Saxena, I. M., Kudlicka, K., Okuda, K., and Brown Jr, R. M. (1994) Characterization of genes in the cellulose-synthesizing operon (acs operon) of *Acetobacter xylinum*: Implications for cellulose crystallization. J. Bacteriol. 176, 5735–5752

8. Standal, R., Iversen, T. G., Coucheron, D. H., Fjaervik, E., Blatny, J. M., and Valla, S. (1994) A new gene required for cellulose production and a gene encoding cellulolytic activity in Acetobacter xylinum are colocalized with the bcs operon. J. Bacteriol. 176, 665–672

9. Kawano, S., Tajima, K., Kono, H., Erata, T., Munekata, M., and Takai, M. (2002) Effects of endogenous endo-β-1,4-glucanase on cellulose biosynthesis in *Acetobacter xylinum* ATCC23769. J. Biosci. Bioeng. 94, 275–281

10. Tajima, K., Nakajima, K., Yamashita, H., Shiba, T., Munekata, M., and Takai, M. (2001) Cloning and sequencing of the beta-glucosidase gene from *Acetobacter xylinum* ATCC 23769. DNA Res. 8, 263–269

11. Tonouchi, N., Tahara, N., Kojima, Y., Nakai, T., Sakai, F., Hayashi, T., Tsuchida, T., and Yoshinaga, F. (1997) A beta-glucosidase gene downstream of the cellulose synthase operon in cellulose-producing acetobacter. Biosci. Biotechnol. Biochem. 61, 1789–1790

12. McNamara, J. T., Morgan, J. L. W., and Zimmer, J. (2015) A Molecular Description of Cellulose Biosynthesis. Annu. Rev. Biochem. 84, 895–921

13. Kumagai, A., Mizuno, M., Kato, N., Nozaki, K., Togawa, E., Yamanaka, S., Okuda, K., Saxena, I. M., and Amano, Y. (2011) Ultrafine Cellulose Fibers Produced by *Asaia bogorensis*, an Acetic Acid Bacterium. Biomacromolecules 12, 2815–2821

14. Penttilä, P. A., Imai, T., Capron, M., Mizuno, M., Amano, Y., Schweins, R., and Sugiyama, J. (2018) Multimethod approach to understand the assembly of cellulose fibrils in the biosynthesis of bacterial cellulose. Cellulose

15. Hu, S. Q., Gao, Y. G., Tajima, K., Sunagawa, N., Zhou, Y., Kawano, S., Fujiwara, T., Yoda, T., Shimura, D., Satoh, Y., Munekata, M., Tanaka, I., and Yao, M. (2010) Structure of bacterial cellulose synthase subunit D octamer with four inner passageways. Proc. Natl. Acad. Sci. U.S.A. 107, 17957–17961

16. Sunagawa, N., Fujiwara, T., Yoda, T., Kawano, S., Satoh, Y., Yao, M., Tajima, K., and Dairi, T. (2013) Cellulose complementing factor (Ccp) is a new member of the cellulose synthase complex (terminal complex) in Acetobacter xylinum. J. Biosci. Bioeng. 115, 607–612

17. Omadjela, O., Narahari, A., Strumillo, J., Melida, H., Mazur, O., Bulone, V., and Zimmer, J. (2013) BcsA and BcsB form the catalytically active core of bacterial cellulose synthase sufficient for in vitro cellulose synthesis. Proc. Natl. Acad. Sci. U.S.A.110, 17856–17861

18. Imai, T., Sun, S.-j., Horikawa, Y., Wada, M., and Sugiyama, J. (2014) Functional reconstitution of cellulose synthase in *Escherichia coli*. Biomacromolecules 15, 4206–4213

19. Du, J., Vepachedu, V., Cho, S. H., Kumar, M., and Nixon, B. T. (2016) Structure of the cellulose synthase complex of *Gluconacetobacter hansenii* at 23.4 A resolution. PLoS ONE 11, e0155886

20. Sun, S.-j., Imai, T., Sugiyama, J., and Kimura, S. (2017) CesA protein is included in the terminal complex of Acetobacter. Cellulose 24, 2017–2027

21. Sun, S.-j., Horikawa, Y., Wada, M., Sugiyama, J., and Imai, T. (2016) Site-directed mutagenesis of bacterial cellulose synthase highlights sulfur–arene interaction as key to catalysis. Carbohydr. Res. 434, 99–106

22. Acheson, J. F., Ho, R. Y., Goularte, N. F., Cegelski, L., and Zimmer, J. (2021) Molecular organization of the *E. coli* cellulose synthase macrocomplex. Nat. Struct. Mol. Biol. 28, 310–318

23. Santos, C. R., Paiva, J. H., Sforça, M. L., Neves, J. L., Navarro, R. Z., Cota, J., Akao, P. K., Hoffmam, Z. B., Meza, A. N., Smetana, J. H., Nogueira, M. L., Polikarpov, I., Xavier-Neto, J., Squina, F. M., Ward, R. J., Ruller, R., Zeri, A. C., and Murakami, M. T. (2012) Dissecting structure-function-stability relationships of a thermostable GH5-CBM3 cellulase from *Bacillus subtilis* 168. Biochem. J. 441, 95–104

24. Acheson, J. F., Derewenda, Z. S., and Zimmer, J. (2019) Architecture of the Cellulose Synthase Outer Membrane Channel and Its Association with the Periplasmic TPR Domain. Structure 27, 1855–1861.e3

25. Nojima, S., Fujishima, A., Kato, K., Ohuchi, K., Shimizu, N., Yonezawa, K., Tajima, K., and Yao, M. (2017) Crystal structure of the flexible tandem repeat domain of bacterial cellulose synthesis subunit. Sci. Rep. 7, 13018

26. Perez-Riba, A., and Itzhaki, L. S. (2019) The tetratricopeptide-repeat motif is a versatile platform that enables diverse modes of molecular recognition. Curr. Opin. Struct. Biol. 54, 43–49

27. Abidi, W., Zouhir, S., Caleechurn, M., Roche, S., and Krasteva, P. V. (2021) Architecture and regulation of an enterobacterial cellulose secretion system. Sci. Adv. 7, eabd8049

28. Uto, T., Ikeda, Y., Sunagawa, N., Tajima, K., Yao, M., and Yui, T. (2021) Molecular Dynamics Simulation of Cellulose Synthase Subunit D Octamer with Cellulose Chains from Acetic Acid Bacteria: Insight into Dynamic Behaviors and Thermodynamics on Substrate Recognition. J.Chem.Theory Comput. 17, 488–496

29. Toyosaki, H., Naritomi, T., Seto, A., Matsuoka, M., Tsuchida, T., and Yoshinaga, F. (1995) Screening of Bacterial Cellulose-producing Acetobacter Strains Suitable for Agitated Culture. Biosci. Biotechnol. Biochem. 59, 1498–1502

30. Nakai, T., Moriya, A., Tonouchi, N., Tsuchida, T., Yoshinaga, F., Horinouchi, S., Sone, Y., Mori, H., Sakai, F., and Hayashi, T. (1998) Control of expression by the cellulose synthase (bcsA) promoter region from *Acetobacter xylinum* BPR 2001. Gene 213, 93–100

31. Rao, F., Pasunooti, S., Ng, Y., Zhuo, W., Lim, L., Liu, A. W., and Liang, Z. X. (2009) Enzymatic synthesis of c-di-GMP using a thermophilic diguanylate cyclase. Anal. Biochem. 389, 138–142

32. Crameri, A., Whitehorn, E. A., Tate, E., and Stemmer, W. P. C. (1996) Improved green fluorescent protein by molecular evolution using DNA shuffling. Nat. Biotechnol. 14, 315–319

33. https://www.gene.mie-u.ac.jp/Protocol/Original/Transform-Comp-Cell-Freeze.html, the latest access date: April 7 2022

34. Updegraff, D. M. (1969) Semimicro determination of cellulose inbiological materials. Anal. Biochem. 32, 420–424

35. Stahlberg, J., Johansson, G., and Pettersson, G. (1993) *Trichoderma reesei* has no true exo-cellulase: All intact and truncated cellulase produce new reducing end groups on cellulose. Biochim. Biophys. Acta-Gen. Subj. 1157, 107–113

36. Studier, F. W. (2005) Protein production by auto-induction in high density shaking cultures. Protein Expr. Purif. 41, 207–234

37. Hashimoto, A., Shimono, K., Horikawa, Y., Ichikawa, T., Wada, M., Imai, T., and Sugiyama, J. (2011) Extraction of cellulose-synthesizing activity of *Gluconacetobacter xylinus* by alkylmaltoside. Carbohydr. Res. 346, 2760–2768

