## Supplementary information for "Dynamic interaction of BcsD subunit in type I bacterial cellulose synthase"

**Experimental procedures**

**Deletion of the *bcsA* gene in *Escherichia coli* BL21**

*Escherichia coli* strain BL21 and the Red/ET recombination system were purchased from New England Laboratory Inc. (MA, USA) and Gene Bridges GmbH (Heidelberg, Germany), respectively. The primers for PCR were synthesized by Fasmac Co. Ltd. (Tokyo, Japan) and were used to amplify the DNA fragment for gene knockout by homologous recombination via the Red/ET system. The DNA fragment consisting of the neomycin-resistance gene cassette (FRT-PGK-gb2-neo-FRT) and homology arm sequences corresponding to the regions flanking the *bcsA* gene of *E. coli* BL21 was prepared by PCR with PrimeSTAR GXL (Takara Co. Ltd., Shiga, Japan). The homology arm sequences were designed based on the complete genome sequence of *E. coli* BL21 (CP010816.1).

5′- CGGCTGAAGAGATACTGACGCTGGCGAACTGGTGCCTGTTGAACTATTCC-3′ (upstream of *bcsA* in sense) and 5′- ACAGCTGGCTCAGCATTGATCAGTGGTTGCGTTGCTGGCGTCGCCTGCGT-3′ (downstream of *bcsA* or the beginning of *bcsB* in antisense). The primer sequences used for this PCR are listed in Table S1.

First, *E. coli* BL21 cells were cultured in LB medium at 37°C until the OD_600_ reached 0.5. The cultured cells were pelleted by centrifugation (11,000 rpm for 30 s at 2°C), washed thrice with sterile 10% glycerol in water, and resuspended in the same solvent. The cell suspension was mixed with 10 ng of pRedET DNA and placed in a 1 mm-gap electroporation cuvette (Nepagene Co. Ltd., Chiba, Japan) for electroporation at a voltage of 1350 V in the Eppendorf Eporator (Eppendorf Corp., Hamburg, Germany). LB medium (1 mL) was immediately added to the cuvette and then recovered into polypropylene microtubes for culturing at 30°C for 1.5 h. Transformed *E. coli* BL21 cells were selected on LB agar plates containing tetracycline (the first transformation).

The tetracycline-resistant colonies from the first transformation were cultured in LB medium containing tetracycline at 30°C. After 20 min, L-arabinose was added to a final concentration of 0.25% (w/v) and the culture was grown until the OD_600_ reached 0.5. The culture flask was placed in ice water for 5 min to cool the cells. The cells were pelleted by centrifugation (11,000 rpm for 30 s at 2°C), washed thrice, and resuspended in 10 % glycerol. The cell suspension was mixed with 800 ng of the neomycin-resistance gene cassette with homology arms described previously, and then placed in a 1 mm-gap electroporation cuvette for electroporation under the conditions same as the first transformation. Electroporated cells were recovered in SOC medium at 37°C for expulsion of the temperature-sensitive pRedET plasmid DNA out of the cell. The transformed *E. coli* BL21 cells were selected on LB agar plates containing kanamycin.

**Construction of the expression vector for the tDGC protein**

The vector used for expression of the tDGC protein in this study was pBAD33et32, a custom-made plasmid. First, we constructed the pBAD202R vector using pBAD202/D-TOPO (Invitrogen Corp., MA, USA.). The TOPO reaction was performed following the manufacturer’s protocol with the following primer pair:

Forward primer: 5′- CACCGAATTCGTCGACGGTACCGCTAGC-3′

Reverse primer: 5′- GCTAGCGGTACCGTCGACGAATTC-3′

The obtained circular plasmid, pBAD202R, was subjected to double digestion with the restriction enzymes MluI and HindIII to obtain the DNA fragment consisting of the BAD promoter, thioredoxin gene (*trx*), and multiple cloning sites. This fragment was ligated into a second plasmid, pBAD33 containing p15A *ori* (supplied by NBRC, NITE, Japan), which was also digested with MluI and HindIII, using the Takara Ligation kit v. 2.1 (TakaraBio Inc., Japan). This vector was named pBAD33/202R and is available for expression of the protein of interest with thioredoxin fused to its N-terminus.

One of the two NcoI restriction sites in the chloramphenicol-resistance gene cassette (Cam^R^) in pBAD33/202R was abolished by site-directed mutagenesis using PCR amplification of the whole plasmid DNA with the following primer pair (mutated bases are underlined):

Forward primer: 5′-ccgttttcacgatgggcaaatattatac-3′

Reverse primer: 5′-gtataatatttgcccatcgtgaaaacgg-3′

The resultant plasmid DNA had a single-base mutation (CCATGG to CGATGG) and was named pBAD33/202RII.

The *trx* gene in pBAD33/202RII, a His-patch thioredoxin derived from pBAD202-TOPO, was replaced by *trx* in pET-32a (Merck Corp.., Darmstadt, Germany), which was PCR-amplified using the following primer pair:

Forward primer: 5′-CCATGGGTAGCGATAAAATTATTCACCTG-3′

Reverse primer: 5′-gagctcgtcgacggtacccatatgagctTTGTCGTCGTCGTCGCCAGAACCAGAACCGGCCAG-3′

This PCR amplification added four restriction enzyme recognition sites, NdeI, KpnI, SalI, and SacI at the 3′ end of the *trx* gene. The DNA sequence was verified after cloning the amplicon into pGEM-T Easy (Promega Inc., WI, USA). The DNA region consisting of *trx* and the four cloning sites was cut out with double digestion using the restriction enzymes NcoI and SacI and then ligated into pBAD33/202RII, whose His-patch *trx* region was excised by double digestion using NcoI and SacI. This plasmid, named pBAD33et32, expresses the protein of interest with thioredoxin fused to its N-terminus and contains five multiple cloning sites: NdeI, KpnI, SalI, SacI, and HindIII. Finally, the gene for tDGC was inserted between the NdeI and HindIII restriction sites in pBAD33et32.

**Characterization of BsCel5A**

The cellulose-binding test was performed with 10 μg of the purified BsCel5A protein in 1 mL of 0.1% (w/v) PASC in 1× PBS. After incubation with rotation for 1 h at 4°C, PASC was pelleted down by centrifugation (10,000 × *g* for 10 min) and washed thrice with 1× PBS. One hundred microliter of 2× SDS-PAGE sample buffer was added to the pellet, and the supernatant was analyzed using a precast 5%−20% polyacrylamide gel (SuperSep Ace).

Next, the enzymatic activity of BsCel5A was tested using several different substrates, namely barley β-glucan, carboxymethyl cellulose, PASC, and Avicel PH-101. The reaction volume was 100 μL, and the concentration of enzyme and substrate was 0.35 μM and 0.1% (w/v), respectively. Two different conditions were tested: (i) 50°C for 15 min in 0.1 M sodium citrate buffer (pH 6.0), or (ii) 4°C for 180 min in 20 mM sodium phosphate buffer (pH 7.5), 300 mM NaCl, 8% glycerol, 5 mM MgCl_2_, and 20 mM imidazole pH 8.0 (wash buffer in metal affinity chromatography for BcsABCD purification without DDM and cellobiose). The former aimed to estimate the approximate full enzymatic activity, while the latter was used to check whether BsCel5A exhibits cellulose-hydrolyzing activity in the BcsABCD purification buffer. The enzymatic reaction was stopped by boiling the mixture for 3 min, and the free reducing sugars in the supernatant were quantified using the Somogyi–Nelson method (1).

Lastly, the substrate specificity of BsCel5A was surveyed using different lengths of cello-oligosaccharides. Each cello-oligosaccharide (DP = 2 ~ 6; Megazyme International Ireland, Wicklow, Ireland) was incubated with 1.5 μM BsCel5A enzyme for 60 min at 50°C in 50 mM sodium citrate buffer (pH 6.0). Next, 10 μL of the reaction solution was spotted on a thin layer chromatography (TLC) plate and developed in TLC solvent (volume ratio 9:6:2 of chloroform: methanol: water), followed by carbohydrate coloration using 10% H_2_SO_4_ in methanol at 120℃.

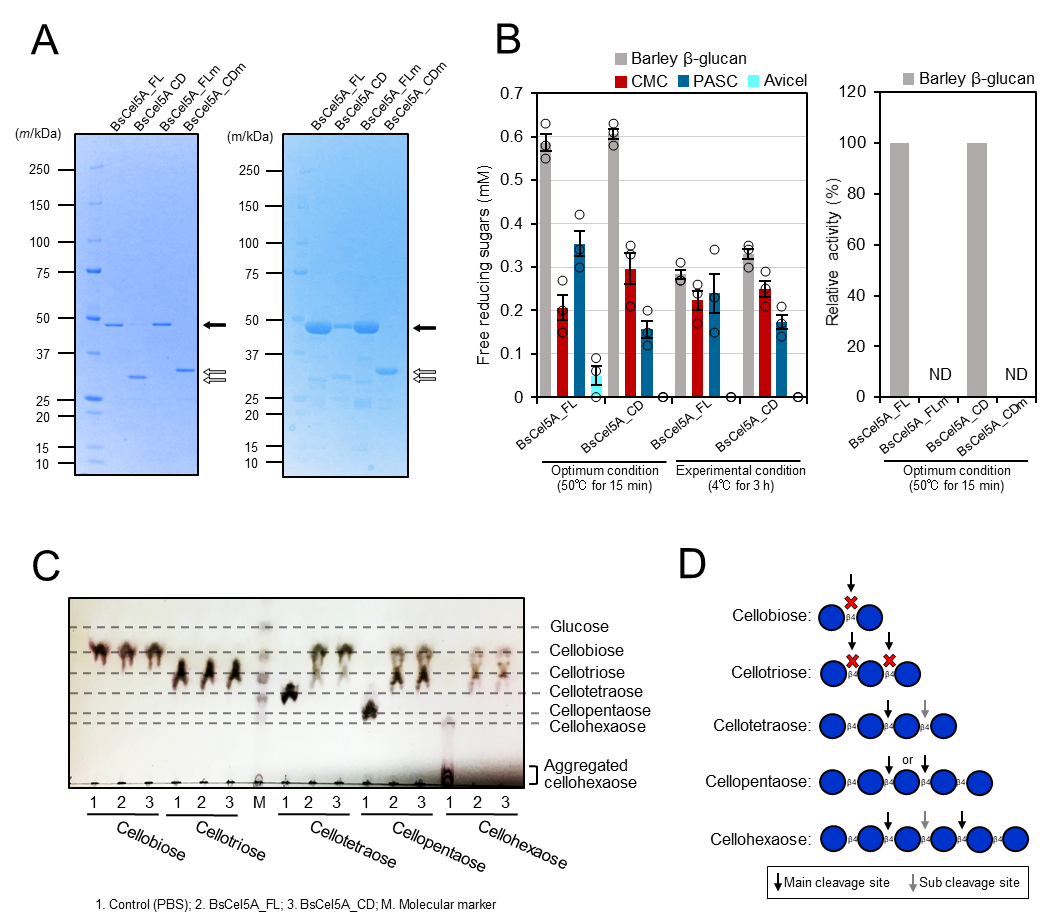

Fig. S1. Characterization of recombinant BsCel5A

**A**, The left panel shows results of the CBB staining of purified BsCel5A, and the right panel shows the results of the cellulose binding activity of purified BsCel5A. Black arrows indicate full-length BsCel5A (BsCel5A_FL) and white arrows indicate the catalytic domain of BsCel5A (BsCel5A_CD). A slight difference in electrophoretic mobility was observed between the wildtype (BsCel5A_CD) and the double mutant E169Q/E257Q (BsCel5A_CDm) due to the amino acid substitutions. **B**, The enzymatic activity of each BsCel5A against various polysaccharide substrates. The enzymatic reactions were carried out under optimal and experimental conditions. The experiments were performed in triplicate (*n* = 3) and are expressed as mean ± SEM. The data values are shown in each plot. **C**, Substrate specificity analysis of each cello-oligosaccharide. **D**, Schematic diagram of the putative cleavage site of BsCel5A for cello-oligosaccharides. The experimental conditions are described in detail in the text.

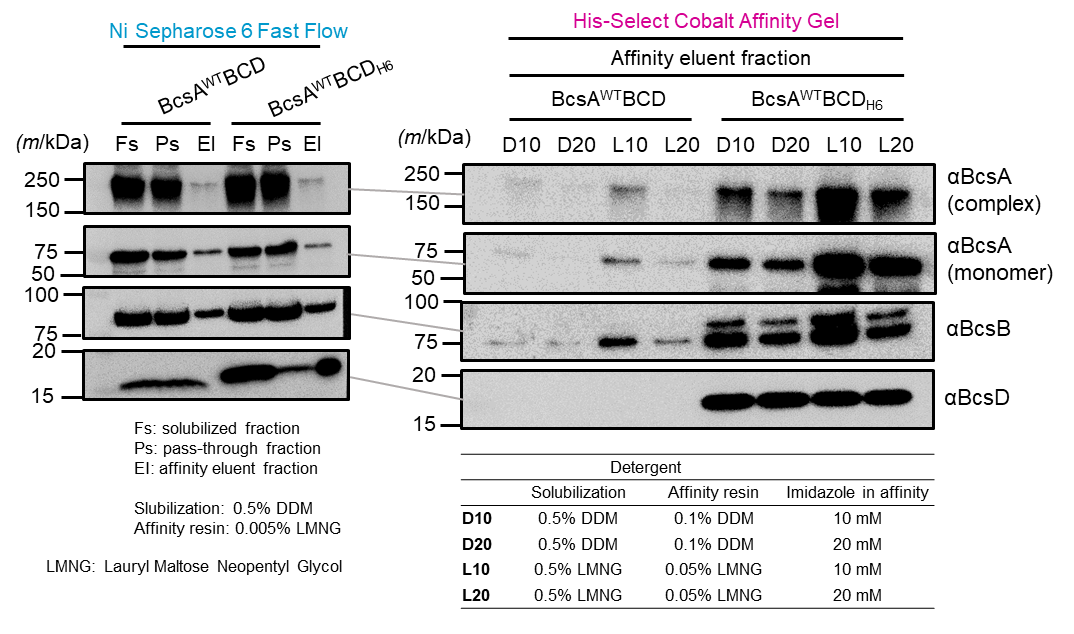

Fig. S2. Selection of affinity resin for BcsABCD co-purification

Recombinant BcsA^WT^BCD_H6_ after IMAC affinity purification was visualized with western blotting using anti-Bcs antibodies. The left and right panels show the results of purification using Ni^2＋^ and Co^2＋^ affinity resins, respectively.

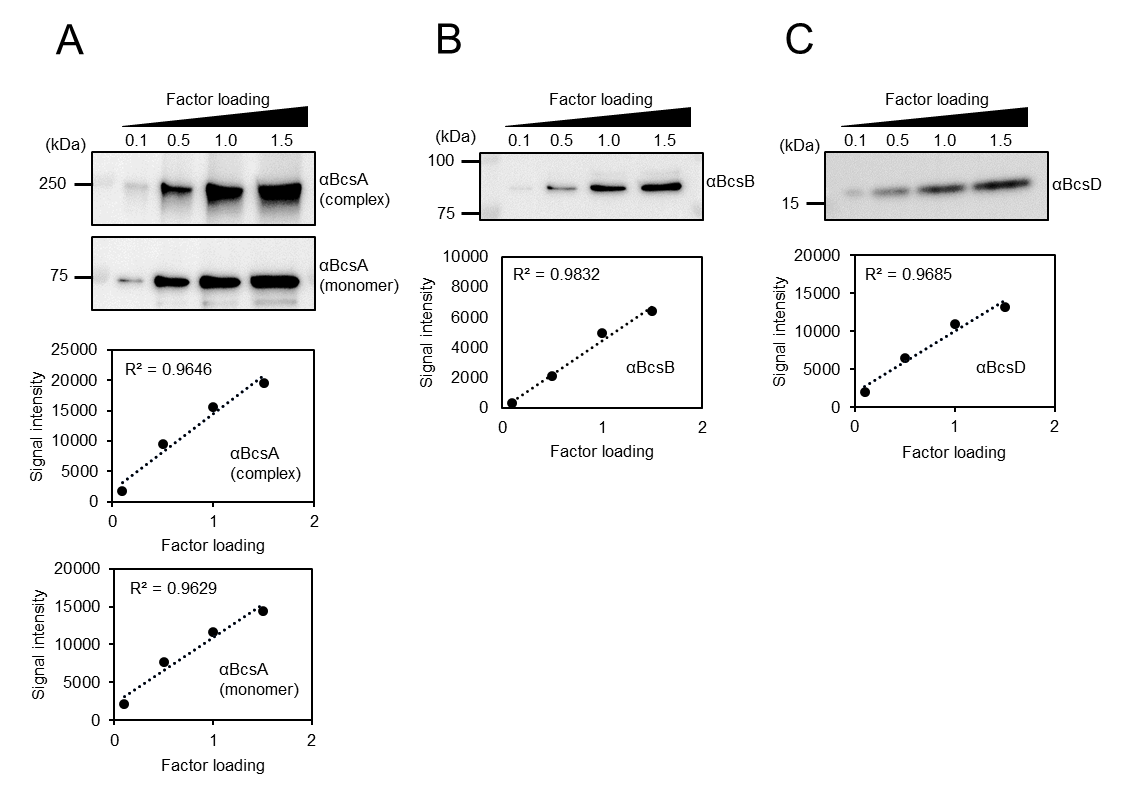

**Fig. S3. Quantitative properties of polyclonal Bcs primary antibodies**

Confirmation of the quantitative nature of the anti-Bcs antibodies. The upper panel shows the western blot image, and the lower panel shows the plot of the signal intensity. **A,** Results of the anti-BcsA antibody. Complex and monomer were split and analyzed separately. **B**, Results of the anti-BcsB antibody. **C**, Results of the anti-BcsD antibody.

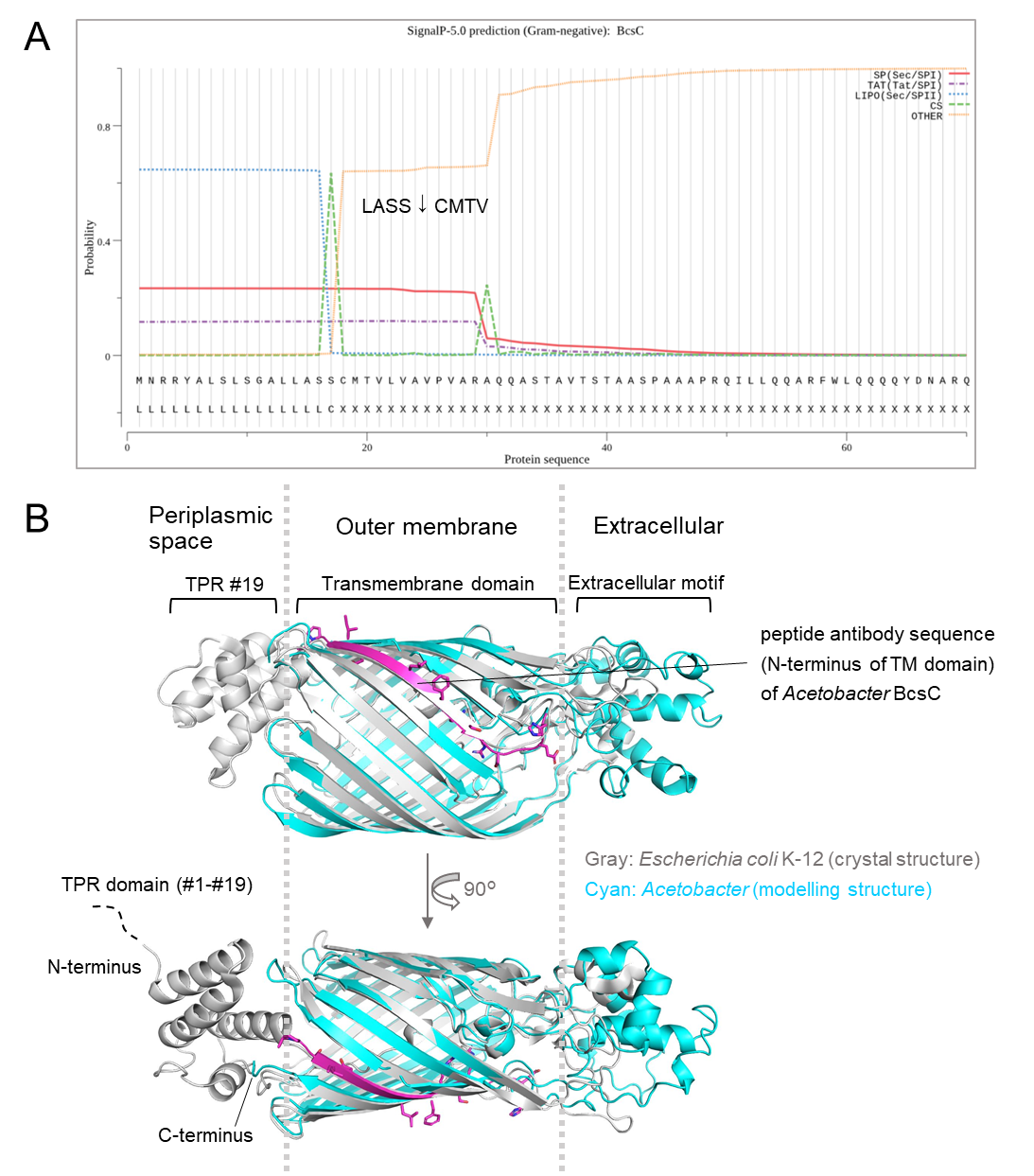

Fig. S4. Predicted structure and signal sequence of BcsC

**A,** The signal sequence of the full-length BcsC amino acid sequence (LC687564) of *Acetobacter* was analyzed using SignalP v. 5.0 (2)

**B**, Superimposition of the ribbon models of *Escherichia coli* K-12 (gray, protein data bank ID: 6TZK) and *Acetobacter* (cyan and magenta). The model structure of *Acetobacter* BcsC was constructed using AlphaFold2 via Google Colab (3).

Table S1. Primer sequences for gene-disruption using the Red/ET recombination system

| Primer name | Sequence |
| --- | --- |
| FR#1 | 5′-ACAGCTGGCTCAGCATTGATCAGTGGTTGCGTTGCTGGCGTCGCCTGCGTAATTAACCCTCACTAAAGGGCG-3′ |
| FR#2 | 5′- CGGCTGAAGAGATACTGACGCTGGCGAACTGGTGCCTGTTGAACTATTCCTAATACGACTCACTATAGGGCTC-3′ |
| #1 | 5′-AAGAGATACTGACGCTGGCGAACTGG-3′ |
| #2 | 5′-TCAGAAGAACTCGTCAAGAAGGC-3′ |
| #3 | 5′-ACAGCTGGCTCAGCATTGATCAGTGG-3′ |

The underlined sequences indicate the homology arm regions, which correspond to the upstream (FR#1) and downstream (FR#2) sequences of the *bcsA* gene of *E. coli* BL21, while the other parts correspond to the sequences at the edge of the FRT-PGK-gb2-neo-FRT gene cassette.

**SI References**

1.	M. Somogyi, Notes on sugar determination. *J Biol. Chem.* **195**, 19-23 (1952).

2.	J. J. Almagro Armenteros *et al.*, SignalP 5.0 improves signal peptide predictions using deep neural networks. *Nat. Biotechnol.* **37**, 420-423 (2019).

3.	M. Mirdita *et al.*, ColabFold - Making protein folding accessible to all. *bioRxiv*, 2 https://doi.org/10.1101/2021.08.15.456425 Deposited February 08, 2022.
